# A Genome Sequence Variant Monitoring Program for Seasonal Influenza A H3N2 and Respiratory Syncytial Virus A using Wastewater-Based Surveillance in Ontario, Canada

**DOI:** 10.1101/2025.07.29.667219

**Authors:** Delaney Nash, Jennifer J. Knapp, Alyssa K. Overton, Yemurayi C. Hungwe, Ria Menon, Jozef I. Nissimov, Trevor C. Charles

## Abstract

Seasonal respiratory viruses, such as the Influenza A virus and the respiratory syncytial virus, are responsible for over a billion infections worldwide each year resulting in a substantial burden on health care systems. Surveillance of these viruses, including their prevalence in communities and their evolution, are essential for informing public health decisions and recommending vaccine formulations and schedules. Typically, these viruses are monitored using clinical samples from patients seeking medical attention. Recently, wastewater-based surveillance (WBS) has been leveraged to understand transmission dynamics and genome evolution of SARS-CoV-2 and seasonal respiratory viruses. To further the utility of WBS we developed and implemented novel tiled-amplicon sequencing assays to identify and track Influenza A virus H3N2 and respiratory syncytial virus A circulating in Southern Ontario, Canada. We also developed virus specific deconvolution tools to estimate the abundance of mixed lineages in wastewater. These assays were able to accurately determine which lineages were circulating in wastewater with high sensitivity and specificity. If implemented in regular surveillance programs, they could be used to inform real-time public health decisions and determine potential disease surge with impact on emergency room visits and hospitalization, as well as track which emerging strains will become predominant in the future and determine which strains should be the focus of seasonal vaccines.

## Introduction

Seasonal respiratory viruses can be highly contagious and typically circulate in the Northern hemisphere from October to April, and in the Southern hemisphere from February to September (1,2). Two seasonal respiratory viruses which cause a large number of yearly infections and substantial health care burden are the Influenza A virus (IAV) and respiratory syncytial virus (RSV) (3–6). These viruses can lead to serious illness and death particularly in vulnerable populations, including children, elderly adults, immunocompromised individuals, and those with comorbidities (5,7). Annually, there are an estimated one billion Influenza infections globally, resulting in approximately 290,000 and 650,000 deaths with the vast majority of these in children under five years old (8). RSV is responsible for an estimated 3.6 million hospitalizations and approximately 100,000 deaths in children under five years old, annually, moreover in 2019 there were an estimated 33 million global RSV-associated acute lower respiratory infection episodes in children under five (9,10). Broader health consequences following an infection can include chronic illness, increased susceptibility to infections, cardiovascular events, and poor pregnancy outcomes, all of which have negative impacts on quality of life and longevity (11–15). Infections can also cause financial strains for individuals who are unable to work due to illness or the need to care for family members (16,17). Additionally, increased hospitalizations and healthcare encounters due to seasonal illness can place a large burden on healthcare and economic systems (12,18–20). In the USA, the healthcare costs attributed to Influenza A and B infections over eight seasons is estimated to be between $2.0-$5.8 billion (20), while estimates of the direct medical cost of RSV infections in the USA range from $1.52-$2.99 billion annually for adults over 60 years of age (21). Routine monitoring along with public awareness campaigns and vaccination of both IAV and RSV is crucial to preventing the spread of these viruses and reducing the burden they impose.

The most common human-infecting IAV subtypes are H3N2 and H1N1(pdm09), with the subtype causing the most infections changing from year-to-year. H3N2-dominated seasons have higher mortality and hospitalization rates in older adults as compared to H1N1(pdm09) or Influenza type B (22). Additionally, H3N2 is known to have a lower vaccine effectiveness and higher immune evasion (22). In the 2022-2023 winter season, the subtype H3N2 was found to be the most prevalent IAV subtype in Canada, making up 89% of IAV subtyped laboratory tests (23), whilst in the following 2023-2024 season, the H1N1(pdm09) subtype was the most prevalent IAV in Canada, making up 73% of IAV laboratory tests with subtype information (24). The IAV H3N2 subtype can be further divided into 35 clades or lineages which are tracked and visualized with NextStrain (https://nextstrain.org/seasonal-flu/h3n2/ha/12y).

Two RSV subtypes, A and B, infect humans and co-circulate during infection seasons (25,26). There are conflicting reports of which subtype causes more severe illness and outcome severity. However, evidence suggests RSV A infections are more prevalent and severe in preterm infants and children with comorbidities (25,26). Definitions at the subgroup level of RSV A and B have been difficult to discern and have been debated within the scientific community (27–29). However, recent work by the RSV Genotyping Consensus Consortium has defined 24 distinct lineages for RSV A (29) which can be visualized with NextStrain (https://nextstrain.org/rsv/a/genome).

Understanding which viral lineages are circulating is important for understanding the disease severity caused by subtypes/lineages, tracking the spatial and temporal spread of viruses, and the effectiveness of vaccines and other treatments (30,31). Traditionally, respiratory virus outbreaks, including those caused by IAV and RSV, have been tracked by testing clinical samples taken from individuals seeking medical treatment. Clinical samples are first tested for viruses using point-of-care diagnostics or reverse-transcription PCR (RT-PCR) analysis, then a subset of positive samples are sequenced to determine the specific subtype or lineage present (23,32–36). However, since the onset of the SARS-CoV-2 pandemic in 2020, wastewater-based surveillance (WBS) has been used alongside clinical testing to track the spread of respiratory viruses in communities (30,37–40). WBS has provided an early warning system for SARS-CoV-2 and other viral outbreaks and has been used to determine which viral subtypes/lineages are circulating in specific areas (38,41,42). WBS offers an attractive additional tool for use alongside clinical testing because it is cost-effective, does not require individuals to seek medical care or report symptoms, and can capture data from asymptomatic or unreported infections in an unbiased and anonymous manner (42–44). Moreover, WBS can inform preventative public health measures which can reduce the health and economic burden imposed by seasonal respiratory viruses (38,42–45).

WBS relies on several molecular biology techniques to fully understand the spread and load of viruses in wastewater (WW), including various qPCR and sequencing techniques (30,38,42). Reverse transcription quantitative PCR (RT-qPCR) methods have been highly successful in tracking the viral load of SARS-CoV-2 and other viruses in wastewater and can determine levels of infection and illness circulating in communities in real time (37,42,45,46). RT-qPCR can also be used to detect and quantify specific viral variants or subtypes in wastewater if they have distinct mutations that distinguish them from other variants/subtypes. This requires the design and testing of specific primers/probe combinations which target differentiating mutations between subtypes (47–49). Thus, subtype monitoring requires a suite of primer/probe combinations to capture subtype diversity, and also demands the frequent development and testing of new primer/probe combinations to detect emerging lineages. Furthermore, virus subtypes such as Influenza A virus H3N2 and RSV A can contain many distinct clades which are rarely discernable using an RT-qPCR approach (40,50).

Sequencing viruses in wastewater offers an attractive alternative to RT-qPCR; viral genomes can be readily amplified using a set of PCR primers inclusive to all lineages of a single virus or virus subtype, with the flexibility to modify the primers when multiple mutations arise in primer binding sites that reduce binding efficiency (51–53). Moreover, sequencing can capture the complete genome sequence, which provides more genomic information and insight into the evolution of virus genomes (54–56). A sequencing approach, dubbed tiled-amplicon sequencing or the ARTIC method, has become a popular PCR-based method to specifically enrich the entire genome of a target virus, and has been widely applied in WBS to detect and characterize variants of SARS-CoV-2 (42,43,51–53).

Since 2020, WBS using a combination of RT-qPCR analysis and targeted sequencing with the ARTIC method has been a powerful public health tool to monitor the viral load and lineages of SARS-CoV-2 in communities. WBS is now expanding to detect other viruses including IAV and RSV (37,39,42,57–60). In the past few years, many studies have been published which monitor the viral load of IAV and RSV in wastewater using RT-qPCR and digital droplet PCR, demonstrating a widespread interest in monitoring these pathogens in wastewater (37,39,46,48,49,57,59–75). However, relatively few studies have utilized genomic sequencing to monitor and evaluate the clade-level variation and genomic evolution of these viruses in wastewater (40,50,54,55,76–80). Among the studies that have embarked on genomic sequencing, some have only utilized a small portion of one genome segment, typically the hemagglutinin (HA) segment, thereby limiting the number of clade specific mutations that can be detected and used for genotyping (50,76,77,80). Other approaches such as a bait-capture (79) and shotgun RNA-Seq (78) did not produce adequate sequencing depth for clade determination. A few studies have used whole genome IAV and RSV targeted sequencing on wastewater samples; however, these studies employed a genome consensus based strategy using clade reference genomes to determine which clades are present in wastewater (40,55). Generating a consensus sequence by mapping reads to a reference genome produces a single representative sequence which includes the most prevalent nucleotide at each position, thus mutations can be removed from data and therefore missed during variant calling which is particularly problematic for identifying rare and low-abundance variants in circulation. Additionally, the abundance of lineages in wastewater cannot be estimated with this strategy. Comparatively, deconvolution methods, with tools such as Freyja (81), observe the prevalence of mutations at clade-defining nucleotide positions and use statistical models to estimate the frequency of mixed-lineages in a sample (82–84). To expand the utility of WBS, we developed novel tiled-amplicon sequencing assays and used them to monitor Influenza A H3N2 and RSV A in wastewater. Samples were collected from six wastewater treatment/collection sites in Ontario, Canada, on a weekly or biweekly basis from November 2023 to January 2024 which coincides with the annual seasonal respiratory virus period in Canada. We applied deconvolution methods to determine which clades were present in samples using Freyja (81), and our in-house tool, Alvoc (abundance learning for variants of concern). Alvoc was adapted for IAV H3N2 and RSV A clade determination (85) from our previously designed tool, Alcov (abundance learning of SARS-CoV-2 variants), which was developed for estimating mixed SARS-CoV-2 lineages in wastewater samples (86). Alvoc was modified to generate constellation files from NextStrain phylogenetic trees. These files delineate clade-specific mutations which are used to call and estimate clade abundances. This modified function enables user adaptation of Alvoc for any virus which has a NextStrain build (85).

Existing primer sets and assays have been previously developed for the detection of IAV and RSV. The MB-Tuni primers (87) and the clinical tiled amplicon RSV (CR) primers (88), have been successfully applied to clinical IAV (32,33,35,87) and RSV (88,89) samples, as well as some agricultural samples (90–92). However, their performance on RNA extracted from wastewater influent has not been thoroughly evaluated, with only one study to our knowledge using MB-Tuni primers on RNA extracted from wastewater (40). Given the fundamental differences between clinical and environmental wastewater samples, including viral concentration, genome integrity, and the presence of inhibitory substances, the MB-Tuni and CR assays may not be amenable to environmental wastewater samples. Thus, we compared our tiled-amplicon assays to these previously designed primer sets using wastewater samples spiked with IAV H3N2 and RSV A virions.

The IAV H3N2 genome consists of eight discrete RNA fragments, called PB2, PB1, PA, HA, NP, NA, MP, and NS, with each segment encoding 1-2 genes (93). In contrast, the RSV A genome is a single RNA strand approximately 15225 bp in length that encodes 10 genes and 11 proteins (94). Our assays employ an ARTIC-style primer scheme, wherein overlapping primers are used to generate ∼400 bp amplicons across each genome. Our IAV H3N2 assay utilizes a total of 42 primers which cover 93.04% of the influenza A H3N2 RefSeq reference genomes NC_007366.1 – NC_007373.1. The RSV A assay utilizes a total of 52 primers which covers 98.44% of the Respiratory syncytial virus A reference genome NC_001803.1.

## Methods and Materials

### Sampling Sites and Wastewater Collection

The study includes data from seasonal wastewater samples collected from six collection sites between November 6^th^, 2023 and January 30^th^, 2024. Either 24 h composite influent wastewater or upstream grab-samples were collected from multiple sites in the Regional Municipalities of York, Peel, Waterloo, and Hamilton (Supplementary Table 1) on a weekly basis, however during a few weeks sample collection was paused at certain sites (Supplementary Table 2). An on site autosampler was used to collect either pre-grit or post-grit separation wastewater influent samples at the GE Booth Water Resource Recovery Facility, as well as the Kitchener, Cambridge, and Hamilton wastewater treatment plants. Grab fractions taken over a 24 h period were pooled to create a composite sample, which was mixed thoroughly to resuspend fecal solids. 250 mL of the composite sample was poured into a sterile polypropylene bottle and refrigerated at 4°C. Samples taken from the two York Region collection sites were three pooled grab samples taken five minutes apart and thus represent a 10-minute time period. The samples were placed in an insulated container with multiple ice packs and shipped to our laboratory in Waterloo, ON, Canada.

### Wastewater and Water Virus Spike-in Samples to Test Primer Specificity

To test primers for specificity and efficiency, wastewater and water samples were spiked with heat-inactivated isolates of IAV H3N2 and RSV A. Upon receiving the wastewater samples, the bottle exteriors were sterilized with 70% ethanol and the bottle was inverted to resuspend solids. Two 20 mL fractions were taken from four wastewater samples and placed in 50 mL falcon tubes. Additionally, four 20 mL volumes of nuclease-free water were placed in falcon tubes to act as spike-in and wastewater controls. One replicate of the four wastewater and two water control samples were spiked with 25 μL of Influenza H3N2 heat-inactivated culture fluid isolate Hong Kong/2671/19 (*Zeptometrix*, #0810609CFHI), while the other replicate of four wastewater samples and two water samples was spiked with 25 μL of RSV-A heat-inactivated culture fluid 2/2015 isolate #2 (*Zeptometrix,* #0810474CFHI).

### Wastewater Virus Isolation and RNA Extraction

Viruses in wastewater and controls were captured and isolated using Ceres Nanotrap Microbiome A beads (*Ceres Nanosciences*, #44202) and RNA was extracted using the QIAGEN RNeasy® Mini Kit (*Qiagen*, #74116). The following amendments to the manufacturer’s protocol for the “Nanotrap® Microbiome A;10mL Manual Protocol with QIAamp® Viral RNA Mini Kit” were made (https://www.ceresnano.com/_files/ugd/f7710c_360d4f6c81094e7e84e161c0c32748b2.pdf). For every 20 mL fraction of wastewater or water, 200 µL of the Ceres Nanotrap® Enhancement Reagent 2 (*Ceres Nanosciences,* #10112-10) and 300 µL of the Microbiome A Particles was used to concentrate the sample. Following a 10 mins incubation period, the Microbiome A beads were pelleted using centrifugation at 8,000 x g for 4 min at 4°C. The pellet was resuspended in a small volume of remaining initial sample instead of nuclease-free water and transferred to a 1.5 mL centrifuge tube for ease of use. The supernatant was removed and 700 µL of RLT lysis buffer (*Qiagen*, #74116) with 1% 2-mercaptoethanol was added to the pellet. The sample was vortexed at maximum speed to release the total virus sized particles into solution and then centrifuged at 8,000 x g for 4 min at RT. The supernatant was transferred to a fresh 2.0 mL round bottom tube. The supernatant with the viral analytes was loaded onto an automated QIAGEN QIAcube Connect® (*Qiagen, #9002864*) device using the pre-loaded manufacturer’s RNeasy® Mini Kit protocol (*Qiagen*, #74116) without on-column DNAse digestion. This virus isolation and RNA extraction method was used for both seasonal wastewater samples and samples spiked with heat-inactivated virus particles.

### IAV H3N2 and RSV A Tiled-Amplicon Primer Design

To design IAV H3N2 genome specific primers, Influenza A H3N2 genomes were downloaded from the NCBI influenza database, applying the following filters; Type: A, Host: Human, Country: Any, Segment: 1-8, Subtype: H3N2, Collection date: 2020-01-01 to 2022-07-21. All segments were downloaded as individual files. The following number of sequences were used for each genome segment alignment: PB2 – 399, PB1 – 401, PA – 402, HA – 416, NP – 406, NA – 409, MP – 407, and NS – 398. Individual multiple sequence alignments (MSA) were generated for each set of the eight segments using the MAFTT-7.0 algorithm. A consensus 90 or consensus 95 sequence was generated from each MSA. Finally, PrimalScheme was used to generate a 400 bp tiled amplicon scheme (Supplementary Table 3) from each consensus sequence. The total length of the generated consensus genome was 13629 bp, of which the primer scheme amplifies 12861 bp (i.e., 94.36% of the genome). Primers were also aligned to the IAV H3N2 RefSeq genome segments, accessions NC_007366.1 – NC_007373.1, and covered 93.04% of this reference sequence. Primers in tiled amplicon schemes are not able to capture the ends of genomic fragments, due to the segmented nature of IAV genomes there is less coverage of the IAV H3N2 than the RSV A genome.

To design RSV A genome specific primers, RSV genomes were downloaded from the NCBI Virus database which met the follow filtering requirements: Virus: Human respiratory syncytial virus A, taxid:208893, Sequence Length: Min - 15200 Max - 15230. A total of 123 genomes were downloaded along with the RSV A RefSeq sequence, NC_001803.1. A MSA was created using the MAFTT-7.0 algorithm and a consensus 85 sequence 15319 bp in length was generated. Finally, PrimalScheme was used to generate a 400 bp tiled amplicon scheme (Supplementary Table 4) from the consensus 85 sequence, which covers 97.84% of the consensus RSV A genome. Primers were also aligned to the RefSeq genome, accession NC_001803.1, and covered 98.44% of this reference sequence.

### Primer Pooling

Initially, primers were pooled at equal concentrations, then were used to PCR amplify cDNA synthesized from RNA that was extracted from a water sample which had been spiked with either Influenza A H3N2 or RSV A heat-inactivated fluid cultures. Ceres bead and spin column extraction was used, as described previously. Following amplicon data analysis, as described below, primer concentrations were adjusted to either increase the depth of reads obtained for amplicons with low coverage or to decrease the depth of reads obtained for amplicons with high coverage. The designated primer pool and concentrations for IAV H3N2 primers (Supplementary Table 3) and RSV A primer (Supplementary Table 4) are listed. These adjusted primer concentrations were used for all described tiled-amplicon PCR assays.

### cDNA Synthesis and PCR Amplification of RNA Extracts from Seasonal WW Samples

RNA extract from seasonal wastewater samples were used to synthesize cDNA with the LunaScript RT SuperMix kit (*New England Biolabs*, #E3010L). Briefly, 20 μL of RNA elution, 8 μL of LunaScript RT SuperMix, and 12 μL of sterile nuclease-free were mixed in a PCR tube. Samples were placed in a thermocycler, then incubated at 25°C for 2 min, 65°C for 15 min, 95°C for 1 min, and then held at 12°C. RNA was converted to cDNA which was subsequently amplified using our tiled-amplicon primers. Separate reactions were prepared for IAV H3N2 and RSV A amplification and each primer pool, for a total of four PCRs per sample. Briefly, 12.5 μL of Q5 High-Fidelity 2x Master Mix (*New England Biolabs*, #M0492L), 6 μL of cDNA, 1 μL of either primer pool 1 or 2, and 5.5 μL of nuclease-free water were mixed in a 0.2 mL PCR tube. Tubes were incubated on the thermocycler at 98°C for 30 s, followed by 35 cycles of 95°C for 15 s then 63°C for 5 min, afterwards samples were held at 12°C.

### PCR Amplification Methods for Virus Spike-in Samples

RNA extracts from IAV H3N2 and RSV A spike in control samples were prepared using both a one-step method, wherein cDNA and PCR amplification occur in a single tube, and a two-step method, wherein cDNA synthesis and PCR amplification occur in two separate tubes. cDNA used in all two-step amplifications was prepared as described above.

#### One-step Method

MB-Tuni (MB) Influenza A H3N2 one-step PCR: 1.25 μL One-step enzyme (*Gene Biosystems*, #P611), 12.5 μL HiScript II 2x Mix (*Gene Biosystems*, #P611), 1 μL of 5 μM pooled MB-Tuni forward and reverse primers (87), 5 μL RNA extract, and 5.25 μL nuclease-free water. Thermocycling conditions: 25°C for 2 min, 50°C for 30 min, 95°C for 2 min, 5 cycles of 94°C for 30 s, 45°C for 30 s, and 68°C for 5 min, 31 cycles of 94°C for 30 s, 57°C for 30 s, and 68°C for 5 min, then held at 12°C (87).

Tiled-amplicon (TA) IAV H3N2 one-step PCR: 1.25 μL One-step enzyme (*Gene Biosystems*, #P611), 12.5 μL HiScript II 2x Mix (*Gene Biosystems*, #P611), 1 μL of H3N2 pool 1 or 2 primers, 5 μL RNA extract, and 5.25 μL nuclease-free water. Thermocycling conditions: 25°C for 2 min, 50°C for 30 s, 95°C for 2 min, 35 cycles of 95°C for 30 s, 63°C for 30 s, then 72°C for 3 min and samples held at 12°C.

Clinical RSV (CR) tiled-amplicon one-step PCR: 1.25 μL One-step enzyme (*Gene Biosystems*, #P611), 12.5 μL HiScript II 2x Mix (*Gene Biosystems,* #P611), either 2.55 μL CR primer pool 1 or 2.8 μL CR primer pool 2, 5 μL RNA extract (88), and topped up to 25 μL with nuclease-free water. Thermocycling conditions: 25°C for 2 min, 50°C for 30 min, 94°C for 3 min, 35 cycles of 98°C for 10 s, 55°C for 10 s, and 72°C for 3 min, then 72°C for 3 min and held at 12°C (88).

Tiled-amplicon RSV A one-step PCR: 1.25 μL One-step enzyme (*Gene Biosystems*, #P611), 12.5 μL HiScript II 2x Mix (*Gene Biosystems*, #P611), 1 μL of RSV A pool 1 or 2 primers, 5 μL RNA extract, and 5.25 μL nuclease-free water. Thermocycling conditions: 25°C for 2 min, 50°C for 30 min, 94°C for 3 min, 35 cycles of 98°C for 10 s, 55°C for 10 s, and 72°C for 3 min, then 72°C for 3 min and held at 12°C.

#### Two-step method

MB-Tuni Influenza A H3N2 two-step PCR: 12.5 μL Q5 2x Master Mix (*New England Biolabs*, #M0492L), 1 μL of 5 μM pooled MB-Tuni forward and reverse primers (87), 6 μL cDNA, and 5.5 μL nuclease-free water. Thermocycling conditions: 98°C for 30 s, 5 cycles of 95°C for 30 s, 45°C for 30 s, and 63°C for 5 min, then 30 cycles of 95°C for 30 s, 57°C for 30 s, 63°C for 5 min, then held at 12°C.

Clinical RSV Tiled-amplicon two-step PCR: 12.5 μL Q5 2x Master Mix (*New England Biolabs*, #M0492L), either 2.55 μL CR primer pool 1 or 2.8 μL CR primer pool 2 (88), 6 μL of cDNA, and topped up to 25 μL with nuclease-free water. Thermocycling conditions: 98°C for 30 s, 35 cycles of 98°C for 30 s, 55°C for 10 s, 63°C for 5 min, then held at 12°C.

The previously described PCRs for seasonal wastewater samples were used for the two-step tiled-amplicon IAV H3N2 and RSV A spike-in samples.

### Library Preparation and Sequencing

Sequencing libraries were prepared from PCR amplified products using the Illumina DNA prep kit (*Illumina*, #20060059) using half of the manufacturer’s recommended volumes. Briefly, PCR products were tagmented using bead-linked transposomes and then indexed using custom Illumina index primers. Samples were pooled and diluted then sequenced on an Illumina NextSeq 2000 using P1 chemistry and 2×300 sequencing.

### Sequencing Read Quality Control, Alignment, and Calculation of Genome Coverage

Raw reads were trimmed and cleaned up using fastp v0.23.2 with default parameters (95,96). Quality controlled reads were aligned to the IAV reference genome EPI_ISL_1563628 or RSV A reference genome EPI_ISL_412866, downloaded from GISAID (February 14^th^, 2024). bwa mem v0.7.12-r118 was used to index reference genomes and align filtered reads (97). samtools v1.13 was used to generate sam and bam files from bwa alignments (98). Genome coverage was calculated from bam files using samtools v1.13 (98).

### Variant Calling with Freyja and Alvoc

Variant calling was performed with Freyja (81) and Alvoc. Briefly, synthetic read data was generated using DWGSIM v0.1.15 (99). To generate synthetic reads, one genome listed from each IAV H3N2 clade in the NextStrain build (https://nextstrain.org/seasonal-flu/h3n2/ha/6y) and RSV A clade in the NextStrain build (https://nextstrain.org/rsv/a/genome/6y) were downloaded from either Gisaid or NCBI virus (March 3^rd^, 2025). DWGSIM v0.1.15 was used to generate 25,000 synthetic reads from each genome without introducing mutations or off-target reads (99).

To call variants, raw reads from seasonal wastewater sequencing libraries and fastq files containing synthetic reads underwent pre-processing steps, first adaptors were removed using cutadapt 5.0 (100). Then bam files were generated using minimap2 2.22 (101) and aligned against the HA reference genome NC_007366.1 or RSV A reference genome NC_001803.1. Bam files and filtered reads were then piped into both the Freyja (81) and Alvoc (85,86) tools set to call IAV H3N2 or RSV A variants. Alvoc only considers mutations at positions that have a read depth greater than 10 bases. For analysis with Freyja, barcode sets specific to each virus were used (https://github.com/andersen-lab/Freyja-barcodes). For analysis with Alvoc, constellations were generated by parsing the NextStrain trees for the HA segment of H3N2 (https://nextstrain.org/seasonal-flu/h3n2/ha/6y) and the RSV A genome (https://nextstrain.org/rsv/a/genome/all-time), mutations present in at least 90% of the nodes belonging to each clade were included in our constellation files. Heatmaps for each virus were generated by both tools for synthetic read data and samples collected at each of the six sampling sites.

### Intraspecific Amplicon Coverage Calculation

Individual fasta files were generated for each amplicon using the primer scheme consensus sequences; the binding sites of primer pairs were used as the start and end site for each amplicon sequence. Filtered reads from seasonal samples were aligned to each amplicon, bwa v0.7.12-r118 was used to index amplicon and align reads (97). samtools v1.13 was used to generate sam and bam files from bwa alignments, then used to calculate genome coverage from bam files (98).

## Results and Discussion

### Comparison of novel and existing IAV H3N2 and RSV A targeted primers

To assess the robustness and performance of our tiled amplicon assays against the previously designed MB-Tuni and clinical RSV (CR) primers, we tested each primer set on water and wastewater samples spiked with heat-inactivated IAV H3N2 and RSV A culture fluid. The MB-Tuni (87) and CR assays (88) utilize a one-step RT-PCR method, wherein both reactions occur in a single tube. Conversely, our assay conditions employ a two-step method wherein cDNA synthesis is followed by PCR with extended low-temperature extension times. For a comprehensive comparison, both one-step and two-step methods were performed with all four primer sets on virus spiked samples. Assay performance was evaluated using the percentage of genome coverage achieved in each sample, as more efficient and reliable amplification of the target virus genome results in higher genomic coverage (Figure 1).

**Figure 1.**
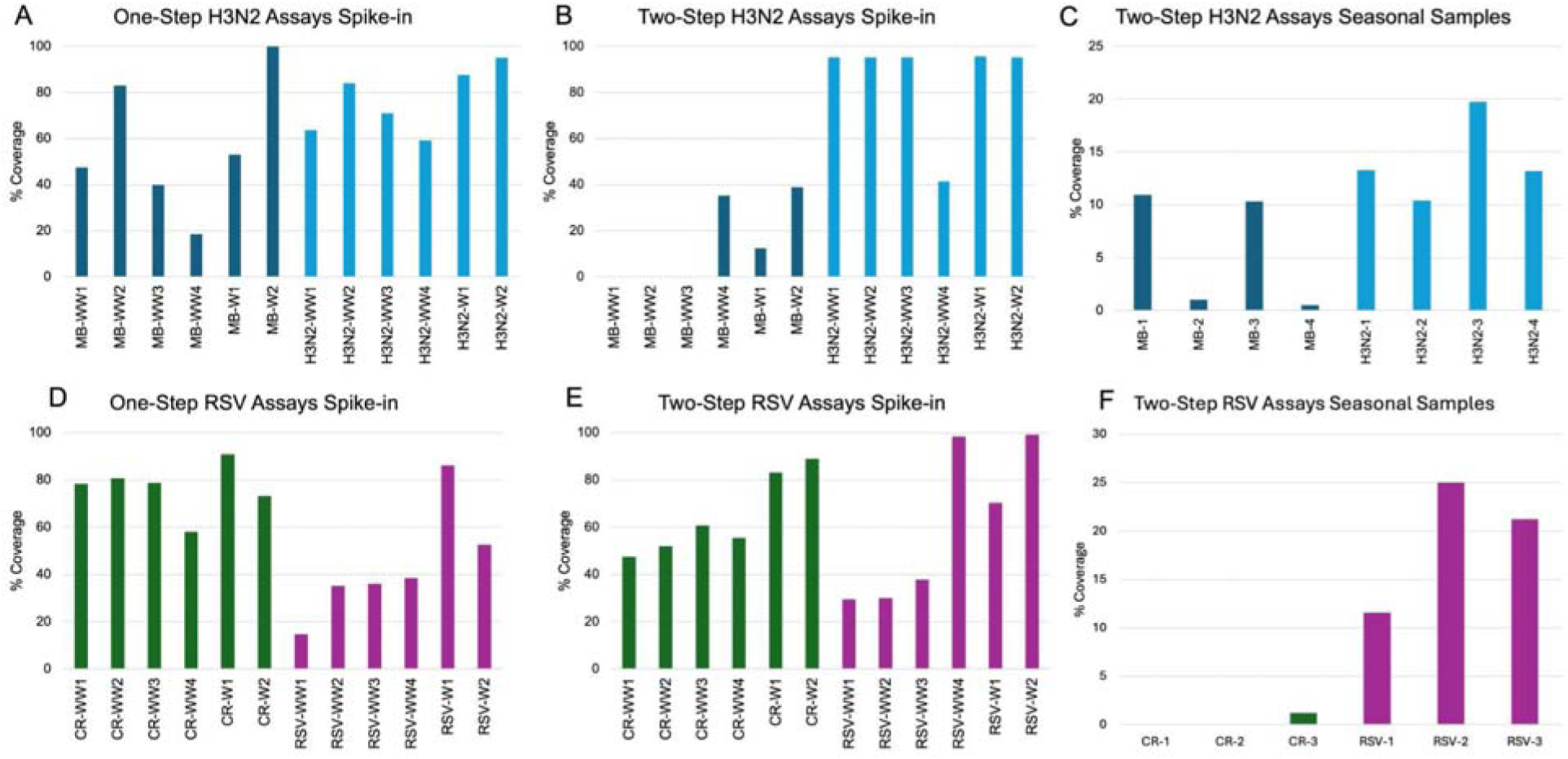
Comparison of Influenza A H3N2 and RSV A amplicon sequencing assays on virus spiked and seasonal wastewater samples. Comparison of IAV H3N2 primer sets, MB-Tuni (MB) and tiled-amplicon H3N2 (H3N2) (A,B,C). Comparison of RSV primer sets, clinical RSV primers (CR) and tiled-amplicon RSV A (RSV) (D,E,F). One-step (A,D) and two-step (B,E) assay efficiency was compared using four wastewater (WW) and two water (W) virus spike-in samples. Seasonal wastewater samples were used to compare two-step assays targeting IAV H3N2 (C) and RSV A (F).

Using a one-step RT-PCR method, the MB-Tuni primers achieved up to 100% genome coverage with virus-spiked water samples. However, they had reduced assay efficiency with virus-spiked wastewater samples, with genome coverage ranging from 19–83% (Figure 1A). In contrast, the TA H3N2 assay achieved up to 95% genome coverage in water and maintained comparatively higher coverage in wastewater samples, ranging from 59–84% (Figure 1A). The superior performance of the TA H3N2 primers in wastewater is likely due to their design, which generates shorter ∼400 bp amplicons. These smaller targets are less susceptible to RNA degradation and inhibitory substances which are common and abundant in wastewater matrices (102). In comparison, the MB-Tuni primers bind at genome termini and amplify across the genome segments which range from ∼800-2300 bp, making them more prone to failure due to RNA segment degradation and the inhibitory PCR conditions (40).

Further evaluation of these assays across one-step and two-step RT-PCR workflows revealed that the MB-Tuni primers performed better with the one-step method, while the TA H3N2 primers showed improved efficiency with the two-step approach (Figure 1A,B). This discrepancy may be attributed to differences in polymerase enzymes and thermocycling conditions between the two workflows, which likely affect primer binding and extension efficiency. Notably, while the MB-Tuni one-step assay’s conditions closely matched those used in the original publication for MB-Tuni primers (87), the two-step method had a modified extension temperature and time. While this modification is more closely aligned with the ARTIC protocol and the two-step TA H3N2 assay, it potentially compromised the MB-Tuni assay performance. However, regardless of the workflow, the TA H3N2 primers consistently produced higher genome coverage in wastewater samples spiked with IAV virions, than the MB-Tuni primers (Figure 1A,B). Additionally, when applied to four unspiked seasonal wastewater samples using the two-step method, the TA H3N2 primers again demonstrated superior genome amplification efficiency across all four samples compared to the MB-Tuni primers (Figure 1C). These findings highlight the TA H3N2 assay’s robustness and suitability for amplifying IAV genomes in complex, inhibitor-rich environmental samples like wastewater.

For RSV detection, the clinical RSV (CR) primers demonstrated strong performance in all spike-in samples, regardless of whether a one-step or two-step RT-PCR approach was used, and consistently outperformed the TA-RSV A primers under high viral load conditions (Figure 1D,E). However, when applied to seasonal wastewater samples without spiked-in virus, the efficiency of the CR assay declined markedly and was surpassed by the TA-RSV A assay (Figure 1F). This discrepancy is likely due to the inherently low concentrations and degradation of viral genomes in environmental wastewater samples. Similar to the MB-Tuni primers used for IAV detection, the CR primers are designed to amplify large genomic regions, with some amplicons extending up to 6,000 bp (88). While effective in controlled spike-in samples and the clinical samples which they were designed for (88), this approach is hindered in real wastewater samples where genomic degradation likely limits the amplification of long fragments (102). In contrast, the TA-RSV A primers amplify shorter ∼400 bp regions, making them less susceptible to the effects of RNA degradation and more reliable for genome recovery in inhibitor-rich environmental wastewater samples.

### Assessment of Genomic Coverage and Comparison to Clinical Percent Positive Tests

As infection rates rise within a community, corresponding increases in viral load within wastewater are expected (64–66). Thus, a robust and efficient PCR assay should yield higher genome coverage and increased sequencing depth as viral load increases in wastewater samples. To assess the performance and epidemiological relevance of our assays, we routinely collected wastewater from six collection sites for three months and used our tiled-amplicon assays on the extracted RNA. We then calculated the percent of genome coverage achieved in each sample and compared these values to the percentage of positive clinical tests reported by the corresponding regional public health units (Figure 2). The six sites we monitored are within the catchments of four public health unit regions: the Cambridge and Kitchener treatment plants in the Region of Waterloo public health and emergency services, GE Booth in the Peel public health unit region, York 1 (Leslie Street), and York 2 (Warden Street) in the York Region public health unit, and the Hamilton collection site in the City of Hamilton public health unit region (Supplementary Table 1, Supplementary Figure 1).

**Figure 2.**
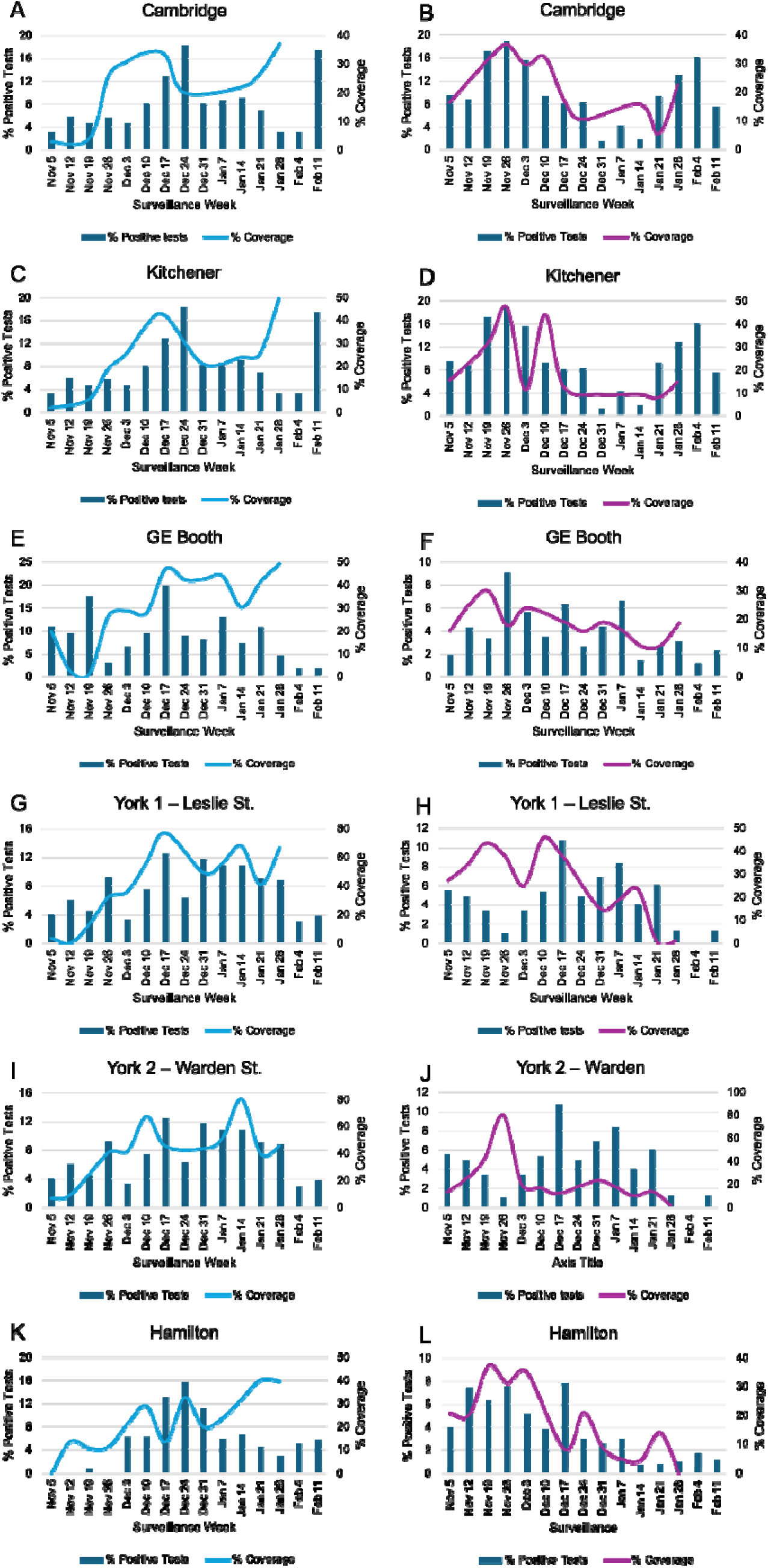
Influenza A H3N2 and RSV A percent of genomic coverage versus percent of weekly positive tests in public health unit regions. IAV H3N2 (a,c,e, g, i, k) and RSV A (b,d,f, h, j, l) genomes in wastewater samples from six sites collected on a weekly or bi-weekly basis were PCR amplified with our tiled-amplicon assays. The percent of genomic coverage (solid lines) was determined for each sample and compared to the percent of positive IAV and RSV tests (bars) reported by public health units in each respective area.

Publicly available surveillance data is reported with the Ontario Respiratory Virus Tool (https://www.publichealthontario.ca/en/Data-and-Analysis/Infectious-Disease/Respiratory-Virus-Tool) and provides weekly percentages of positive tests and number of cases in each public health region. The percent of positive tests was selected over the absolute number of reported cases, as case counts are directly influenced by testing volume, which can fluctuate weekly. By considering the percentage of positive tests, we accounted for changes in the level of testing and obtained a more reliable indicator of infection prevalence within each community. This allowed for a meaningful comparison between wastewater-based genome coverage and infection trends in the population serviced by each respective public health unit (Figure 2).

Generally, of the six sampling sites, we observed that increases in genome coverage generally corresponded with rising percentages of positive tests within the associated public health regions (Figure 2). Notably, peaks in genomic coverage of IAV H3N2 in Kitchener, Cambridge, and York 2, were observed one to three weeks before corresponding increases in clinical positivity rates, suggesting that these assays may provide an early warning of escalating viral transmission within these communities (Figure 2A,C,I). For example, when monitoring IAV H3N2 cases in Cambridge and Kitchener, which are both in the Waterloo region, peaks in genomic coverage were observed in the weeks of December 17th, 2023 and January 28th, 2024, one and two weeks before infection levels peaked in this region (Figure 2A,C). Additionally, peaks in genomic coverage of RSV A in GE Booth, York 1, York 2, and Hamilton, were observed one to three weeks before peaks in clinical positivity rates (Figure 2F,H,J,L). A notable caveat that must be addressed are gaps in testing eligibility and clinical reporting data that may affect the comparisons between genomic coverage and clinical positivity rates. In Ontario, clinical testing is only performed on 4 eligible groups, including: symptomatic children under 18 seen in emergency departments, symptomatic hospitalized patients, symptomatic residents in institutional settings, and the first four symptomatic individuals in an outbreak setting that request testing (https://www.publichealthontario.ca/en/Laboratory-Services/Test-Information-Index/Virus-Respiratory). Thus, clinical testing and positivity rates are biased toward individuals in these groups and do not capture the full scope of infected individuals. Additionally, only a small percentage of IVA and RSV clinical positive tests are subtyped, thus the clinical data includes both IVA subtypes H3N2 and H1N1(pdm09) and RSV A and B subtypes, which could further skew our comparison of clinically positive tests and genomic coverage. Although the current testing data does work as a proxy for the general levels of infection in each public health region, we suspect that if a greater number of individuals were tested and samples were subtyped there may be a better correlation between clinical positivity rates and genomic coverage across all sites and would perhaps alleviate some of the observed discrepancies (Figure 2K).

While genome coverage is not a quantitative proxy for viral load or case numbers, these trends across multiple locations demonstrate the utility of our assays for monitoring virus activity in wastewater and supporting public health surveillance. These findings indicate that wastewater-based genomic surveillance can effectively monitor IAV H3N2 and RSV A infections throughout the respiratory virus season. The increases and decreases in genomic coverage mirror patterns of rising and falling clinically reported cases within the respective collection regions. Thus, these assays offer an accurate real-time assessment of active IAV H3N2 and RSV A cases in municipalities, demonstrating the value and feasibility of monitoring the transmission and prevalence of these seasonal viruses with wastewater-based surveillance. Furthermore, in some instances they have the potential to offer valuable lead-time for public health unit response efforts, however more complete clinical testing of individuals is necessary to properly confirm those trends.

### Evaluation of Synthetic Reads with Freyja and Alvoc

To support the identification and tracking of Influenza A H3N2 (IVA H3N2) and Respiratory Syncytial Virus A (RSV A) variants, we developed a lineage deconvolution tool, Alvoc, to estimate abundance of these viruses in wastewater samples. Similar to its predecessor for SARS-CoV-2 variant deconvolution (86), Alvoc utilizes a normalization approach based on mutation positions characteristic of specific clades and applies a linear regression model via ordinary least squares to estimate variant abundances within a sample (85,86).

Several other variant-calling tools, such as Freyja, have been developed for deconvoluting mixed viral populations. Freyja similarly relies on lists of clade-defining mutations and their genomic positions but employs a constrained weighted approach using least absolute deviations to estimate variant proportions (81). Notably, our original SARS-CoV-2 variant deconvolution tool, Alcov, was previously benchmarked against Freyja and seven additional tools, where both Alcov and Freyja demonstrated strong and comparable performance, accurately resolving SARS-CoV-2 variant mixtures in complex samples (83). Given this, we compared the performance of both Alvoc (i.e., modified Alcov) and Freyja for IAV H3N2 and RSV A detection in wastewater using synthetic read datasets generated from one virus genome per clade.

To evaluate the performance of Alvoc and Freyja for IAV H3N2 and RSV A variant calling, we generated synthetic read data sets from one representative virus genome per clade for each virus and analyzed these datasets using both tools. Both Alvoc and Freyja utilized mutations in the hemagglutinin gene (HA) segment to estimate IAV lineage abundances. In these controlled tests on IAV genomes, the two tools showed comparable performance (Figure 3). Freyja correctly identified 27 out of 31 clades (Figure 3A), while Alvoc identified 26 out of 31 (Figure 3B). However, both tools encountered challenges in differentiating certain closely related clades. For example, clade 3C.2 was frequently misclassified as 3C.2a, and subclades such as 3C.2a and 3C.2a1a were called as the closely related clade 3C.2a1. Additionally, Alvoc misclassified clade 3C.2a1b.2 as 3C.2a1b.2b, demonstrating minor difficulties in resolving fine-scale differences between closely related lineages. The sensitivity, defined as the tools ability to correctly call a clade, and specificity, defined as the tools ability to correctly reject a clade, were determined for each tool. Sensitivity was calculated by dividing the number of true positives (correctly called clades, at <2%) divided by the number of false negatives (correct clades that were not called, at >2%) plus true positives. Specificity was calculated by dividing the number of true negatives (incorrect clades are not called, at >2%) by the number of false positives (incorrect clade was called, at <2%) plus the number of true negatives. The sensitivity and specificity of Freyja IAV calls were 87.10% and 99.57%, respectively, and for Alvoc IAV were 83.87% and 99.46%, respectively. Thus, Freyja is slightly better than Alvoc at correctly calling IAV H3N2 clades, while both are able to correctly reject incorrect clades with a similar frequency.

**Figure 3.**
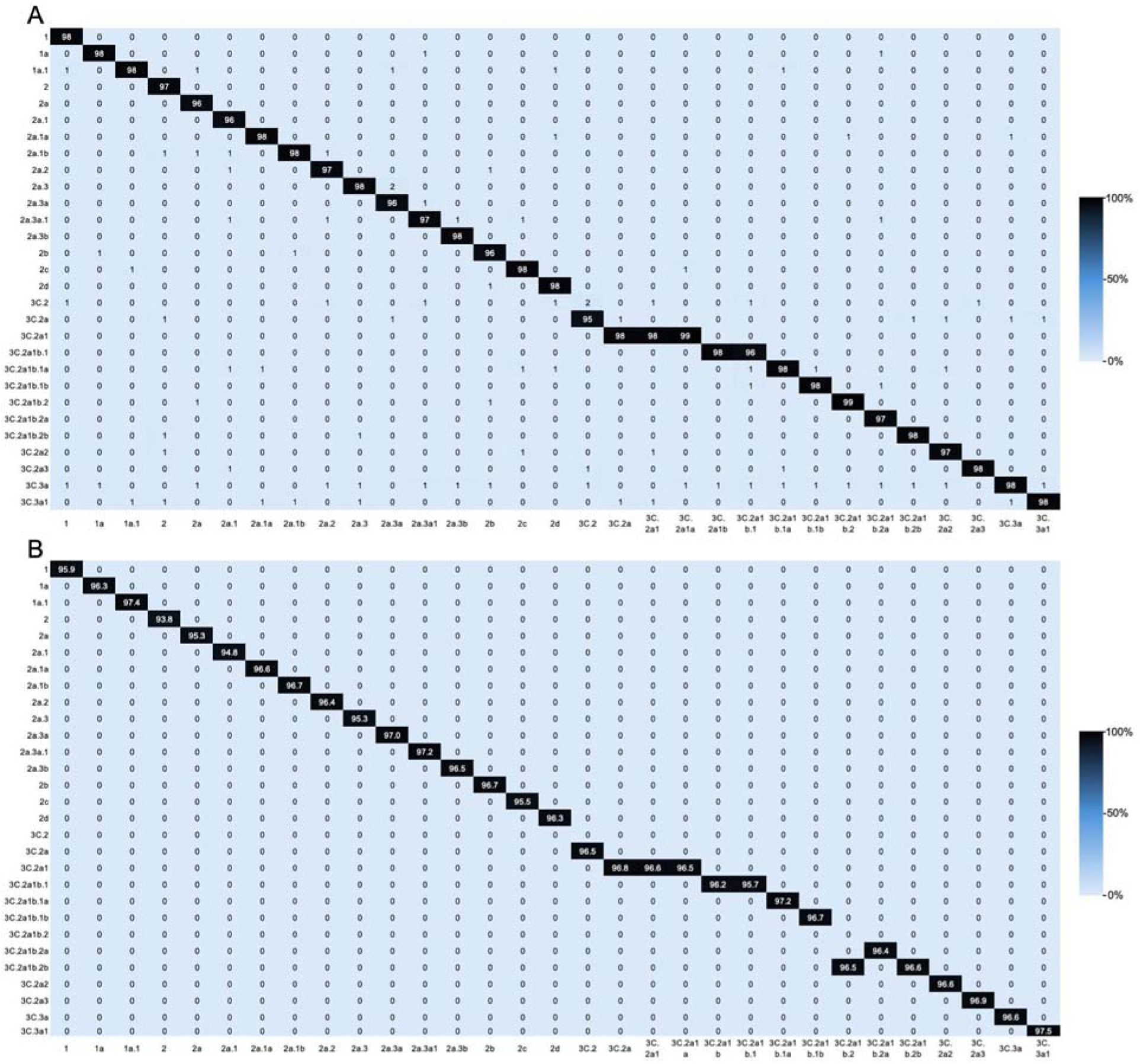
Comparison of Freyja and Alvoc IAV H3N2 variant calling using synthetic read data. Variant calling on synthetic reads, generated from genomes belonging to IAV H3N2 viruses in each clade with DWGSIM, was performed with both Freyja (A) and Alvoc (B). Sample names, i.e. the clade each synthetic read dataset belongs to, is displayed on the horizontal axis, while the clade called for each sample is displayed on the vertical axis. Heatmaps are used to delineate the percentage of reads assigned to each clade. Clades with zero percent calls across all synthetic data sets were excluded from the heatmap.

When analyzing RSV A synthetic read datasets, Alvoc outperformed Freyja (Figure 4). Alvoc successfully identified all RSV A clades (Figure 4B). However, two read data sets were identified as clade mixtures. The A.3.1 read data set was classified as 65.9% A.3.1 and 27% A.D., while A.3.1.1 was classified as 58.7% A.3.1.1 and 32.2% A.D. The misclassifications largely reflected close phylogenetic relationships between these clades which share similar mutations (Figure 4B). In contrast, Freyja correctly called only 23 out of 36 RSV A clades, with frequent misclassifications of subclades being grouped under their parent clade (Figure 4A). For instance, variants A.D.1.4 through A.D.1.8 were consistently miscalled as the parent clade A.D.1 (Figure 4A). This suggests that while both tools are capable of resolving major clades, Alvoc offers greater resolution and specificity when applied to RSV A variant mixtures. The sensitivity and specificity of Freyja RSV A calls were 63.89% and 98.97%, respectively, while for Alvoc RSV A calls were 100% and 99.84%, respectively. Thus, Alvoc was able to correctly identify clades with a much higher frequency then Freyja, while both tools were able to reject incorrect clades with a similar frequency.

**Figure 4.**
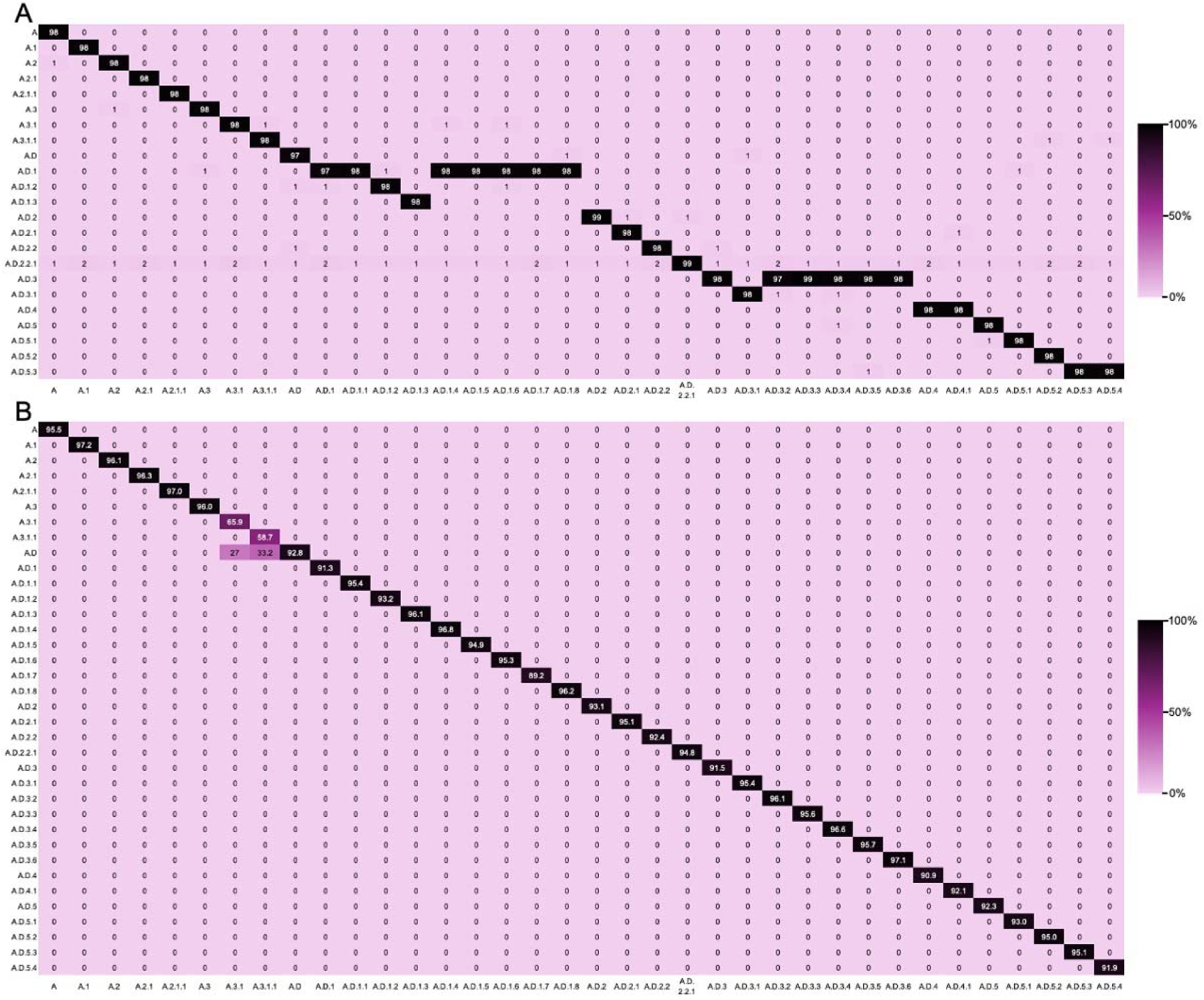
Comparison of Freyja and Alvoc RSV A variant calling using synthetic data. Variant calling on synthetic reads, generated from genomes belonging to RSV A viruses in each clade with DWGSIM, was performed with both Freyja (A) and Alvoc (B). Sample names, i.e. the clade each synthetic read dataset belongs to, is displayed on the horizontal axis, while the clade called for each sample is displayed on the vertical axis. Heatmaps are used to delineate the percentage of reads assigned to each clade. Clades with zero percent calls across all synthetic data sets were excluded from the heatmap.

There are several differences between these tools which likely contribute to the minor discrepancies between variant calls. For one, the statistical method for estimating clade abundances differs, which may result in differing lineage abundance estimates. Most significantly, Freyja and Alvoc utilize different methods and parameters for generating their constellation files which delineate clade-specific mutations; resulting in differences between clade-defining mutations used by each tool. The mutations used by each tool to define clades plays the largest role in what clades are ultimately called, and thus are the most likely source of discrepancies between the variant calling tool output. On top of utilizing different mutations, Freyja’s RSV A constellation files do not define lineages A.D.1.4-A.D.1.8, A.D.3.2-A.D.3.6, or A.D.5.3, which is why these lineages were not correctly called by Freyja (Figure 4). Despite these discrepancies, these results demonstrate that both Alvoc and Freyja are capable of deconvoluting IAV H3N2 variant mixtures in wastewater samples, with Alvoc showing enhanced performance for some closely related RSV A clades. These findings support the feasibility of wastewater-based genomic surveillance for respiratory viruses beyond SARS-CoV-2 and highlight the potential of Alvoc as a valuable tool for public health monitoring of seasonal respiratory virus transmission.

### Application of Freyja and Alvoc for the Assessment of IAV H3N2 and RSV A Clades in Seasonal Wastewater Samples

Freyja and Alvoc were applied to sequencing reads from seasonal wastewater samples to estimate H3N2 lineage abundances. Freyja generated lineage calls for more samples than Alvoc, largely due to differences in minimum read depth thresholds. Freyja operates with a default threshold of zero, thus can include low-coverage positions in its abundance estimates, while Alvoc requires a minimum of 10× depth, limiting its ability to analyze samples with sparse coverage. However, this increased detection capacity in Freyja reduced the certainty of some lineage assignments, and often distributed abundance estimates across multiple clades when coverage was low (Figure 5).

**Figure 5.**
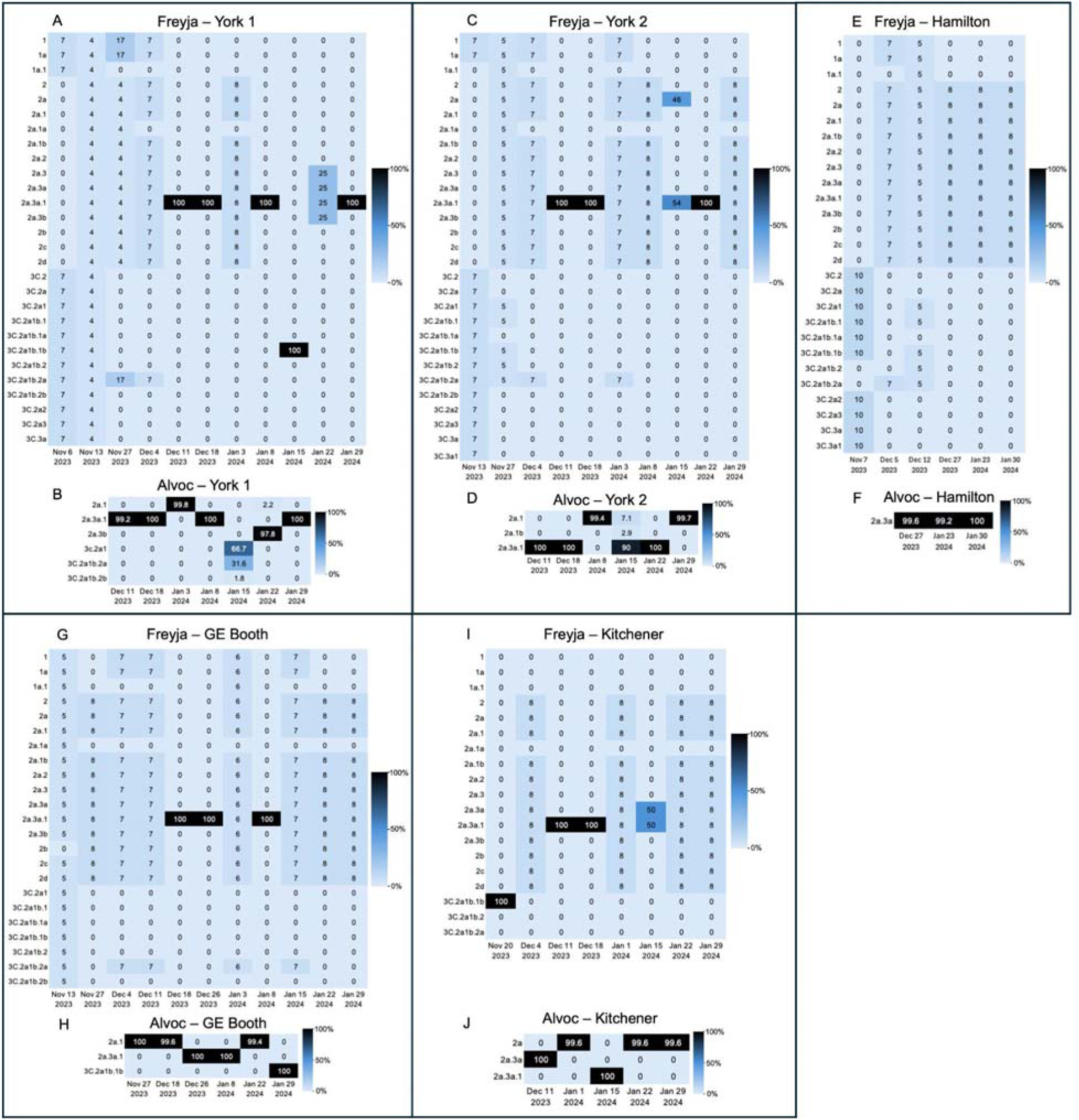
IAV H3N2 variant calls with Freyja and Alvoc from seasonal wastewater samples. Heatmaps display the lineage abundance calls for IAV H3N2 estimated using Freyja (A,C,E,G,I) and Alvoc (B,D,F,H,J). Includes results from seasonal wastewater samples collected at sites in the York Region (A,B,C,D), Hamilton region (E,F) Peel region (G,H), and the Waterloo Region (I,J).

In samples where both tools produced confident calls, the primary H3N2 lineages detected were 2a, 2a.3a, 2a.3a1, and 3C.2a1b.1b (Figure 5). These observations are consistent with global surveillance data. According to the NextStrain H3N2 build, of 143 genomes collected from 42 countries between our sampling period of November 2023 and January 2024, 141 genomes were classified as 2a.3a1, confirming this clade’s predominance internationally. Our Ontario wastewater findings similarly showed widespread detection of 2a.3a1 (Figure 5). In almost all samples where 2a.3a1 was called, both Alvoc and Freyja agreed on these clades assignments. In some samples, for example York1 Jan 3^rd^ and York2 Jan 8^th^, Alvoc called clades 2a, 2a.1, or 2a.3a, instead of the closely related and predominant lineage 2a.3a1. Either, Alvoc had minor difficulties in distinguishing between these closely related clades, or these clades were co-circulating alongside 2a.3a1. The former seems more likely, as 97% of sequenced clinical samples in Canada during this time were assigned to clade 2a.3a1 (103). When Freyja was used to evaluate these same samples, it was unable to determine the specific clade and instead split the lineage abundance call between 12 possible clades (Figure 5). In these samples the HA gene read coverage and/or depth was low, which likely made it difficult for these tools to discern between closely related lineages which share common clade determining mutations. To enhance read recovery, various virion capture and isolation methods, such as PEG precipitation or ultracentrifugation, could be evaluated to determine whether they improve virion isolation, thereby increasing IAV genome concentrations, PCR product yields, and sequencing read counts (57,104,105).

The 2a.3a1 clade (formally 3C.2a1b.2a.2a.3a.1) was initially reported in the United States in 2021, as reported by GISAID, and quickly disseminated worldwide becoming the predominant H3N2 clade in 2023 (104). This clade carries distinct amino acid changes in the HA and neuraminidase (NA) genes as compared to the 2023/2024 WHO-recommended vaccine strains which reduced vaccine protection and efficacy during this season (103,105–107). In Canada, 97% of the sequenced H3N2 infections during the 2023/2024 season belonged to 2a.3a1, and vaccine effectiveness against circulating H3N2 strains was estimated at 40%, indicating moderate protection (103). In the 2024/25 season, the vaccine against the H3N2 strain was updated using a clade 2a.3a1 strain, which increased the vaccine protection to 54%, and was particularly effective against 2a.3a1 strain infections (108). The implementation of a genomic surveillance system like this could provide real-time insight into emerging strains, overcoming delays associated with clinical genotyping of strains. If applied globally, it could help predict which clades are likely to become predominant and better inform vaccine strain selection, improving the vaccines alignment with circulating variants and enhancing their effectiveness (38).

According to the NextStrain RSV A build, multiple RSV A clades were circulating globally during our sampling period, with A.D.5.2 (24% frequency), A.D.1 (20% frequency), and A.D.3 (13% frequency) representing the three most prevalent clades, several additional clades were also detected at lower frequencies. Overall, variant calls generated by both Freyja and Alvoc were broadly consistent, with minor differences in estimated lineage abundances (Figure 6). Discrepancies between tools became more apparent in samples collected toward the end of the RSV season, i.e. the end of December and January, when declining viral RNA concentrations reduced genome recovery, complicating lineage assignment. Notably, in synthetic dataset analyses, Freyja was unable to resolve strains within subclades A.D.3.2–A.D.3.6, instead assigning them to their parent clade A.D.3, whereas Alvoc accurately classified these strains to their respective clades. This pattern was reflected in wastewater samples, where Alvoc detected multiple A.D.3 sub-lineages that Freyja did not resolve (Figure 6). Geographically, differences in clade distributions were observed (Supplementary Figure 1). In the York (York 1 and York 2) and Peel (GE Booth) Public Health region, which are part of the Greater Toronto Area, A.D.5.2 and A.D. 5.1 emerged as the dominant clade. Whereas samples from the Waterloo (Kitchener and Cambridge) and Hamilton regions, which are located closely together, displayed greater clade diversity with no single dominant lineage (Supplementary Figure 1). These findings highlight the regional variation in RSV A transmission dynamics and the ability to capture these dynamics using WBS.

**Figure 6.**
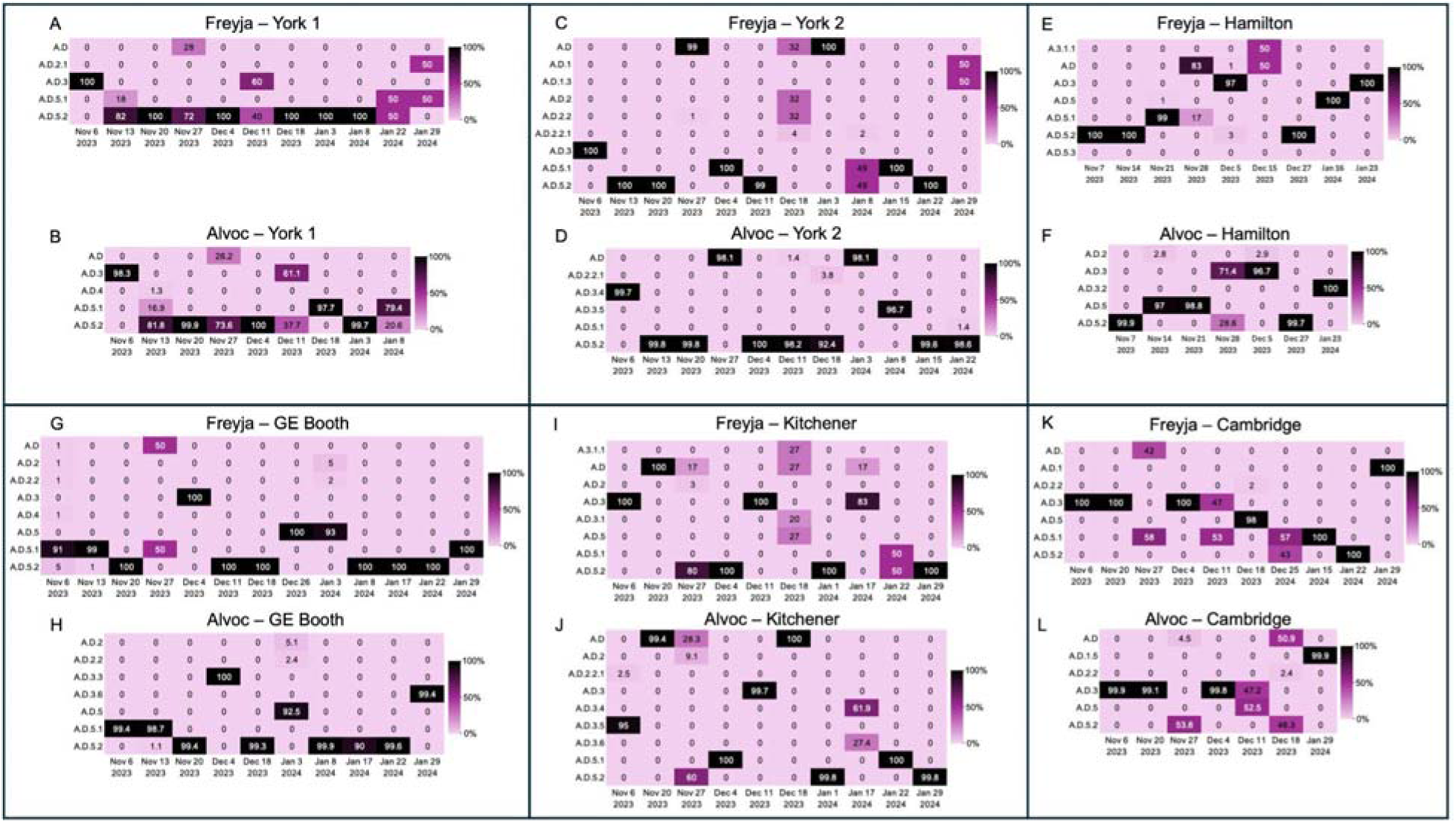
RSV A variant calls with Freyja and Alvoc in wastewater samples. Heatmaps display the lineage abundance calls for RSV A using Freyja (A,C,E,G,I,K) and using Alvoc (B,D,F,H,J,L). Includes results from seasonal wastewater samples collected at sites in the York Region (A,B,C,D), Hamilton region (E,F) Peel region (G,H), and the Waterloo Region (I,J,K,L).

Currently available RSV immunization strategies rely on vaccines incorporating recombinant fusion (F) protein antigens to elicit protective immunity (109), while therapeutic interventions in infected individuals predominantly use monoclonal antibodies targeting the RSV F protein (110,111). Both approaches are susceptible to the effects of viral evolution, particularly mutations in the F protein that can reduce antibody binding and neutralization (111,112). Consequently, ongoing genomic surveillance to track the emergence of potential escape variants is critical for maintaining the effectiveness of these preventive and therapeutic measures (112). Wastewater-based surveillance offers an unbiased, community-level approach for monitoring RSV variant circulation and detecting early signs of immune escape trends.

### Intraspecific Amplicon Analysis of Tiled-Amplicon Sequencing Assays

A total of 67 samples had genomic coverage for both IAV and RSV (Figure 7). Among the IAV H3N2 amplicons, those targeting the PB2 (subunit of the RNA polymerase complex) and NP (nucleoprotein) segments consistently achieved the highest coverage (Figure 7A). Notably, many RT-qPCR assays target the MP (matrix protein) segment (48,75,113–115) and HA segment (47,65,68). However, amplicons in the MP region showed coverage in only 24–29 of the 67 samples, while amplicons 23-26 in the HA region had coverage in 12-17 samples and amplicons 27-28 had coverage in 41 and 45 samples. In contrast, PB2 amplicons 2, 3, 7, and 8 were covered in 56–57 samples, and NP amplicons 31 and 32 in 53 and 55 samples, respectively. This pattern may reflect PCR primer bias, with certain primers exhibiting greater binding efficiency and amplification success. Alternatively, it may indicate differential RNA degradation across genome segments in wastewater, with PB2 and NP regions being more stable. If so, these segments could serve as more reliable targets for RT-qPCR assays of IAV in wastewater.

**Figure 7.**
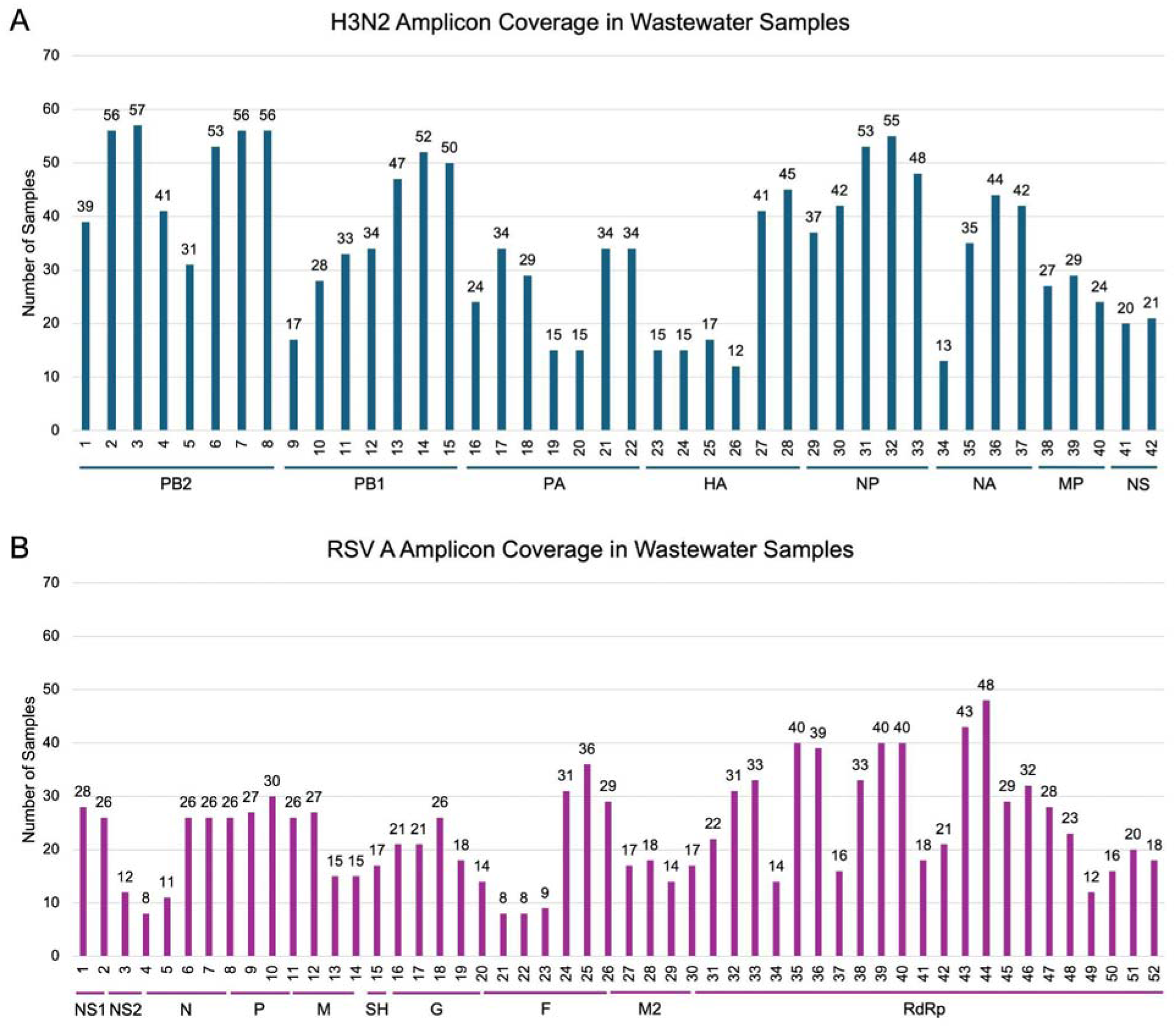
IAV H3N2 and RSV A amplicons with coverage in wastewater samples. The number of samples which contained coverage of IAV H3N2 amplicons (A) and RSV A amplicons (B). A total of 67 samples had genomic coverage of IAV H3N2 and RSV A. The number of samples which had amplicon coverage are labeled above bars. IAV H3N2 assay utilized 42 amplicons, the genome segment each amplicon bound to is represented by blue bars and segment labels (A) (Supplementary Table 5). The RSV A assay utilized 52 amplicons, the gene each amplicon bound to is represented by purple lines and gene labels (B) (Supplementary Table 6).

For RSV A, amplicon 44, which targets the RNA-directed RNA polymerase (RdRp) gene, displayed the highest coverage, present in 48 of the 67 samples (Figure 7B). Some of the currently used RSV A RT-qPCR assays target the N gene (46,50,58,61) and M gene (48). However, these regions had much lower coverage in the N and M genes compared to the RdRp. Similar to our analysis of IAV H3N2 amplicons, this may suggest greater RNA stability in the RdRp region, making it a preferable target for RT-qPCR assays monitoring RSV in environmental samples.

### Future Goals and Assay Optimization

To improve upon our genomic surveillance methods of IAV H3N2 and RSV A, assays and bioinformatic tools could be further optimized. To improve our assays, a variety of experimental modifications regarding virion capture, RNA extraction, and PCR amplification could be evaluated. As previously discussed, virus concentration and isolation techniques could be compared to increase virion yield. Improving RNA extraction from wastewater could also be explored by testing different kits and including a post-extraction DNase treatment step. Variations of our PCR assays could also be tested to optimize and increase the yield of PCR products prior to library preparation. Such variations could include testing of different thermocycling conditions and increasing the concentration of primers that target genomic regions with low yields. Lastly, the feasibility of pooling IAV H3N2 and RSV A primers to reduce the number of PCR reactions and library preparation samples was not explored, and if successful, could significantly reduce the cost of materials and labour required.

In regards to our bioinformatic approaches, improvements to our IAV H3N2 Alvoc deconvolution tool could also be made. Our RSV A deconvolution tool performed exceedingly well, demonstrating high sensitivity and specificity, and was able to call variants in a greater number of samples than our IAV H3N2 tool. This is likely due to its ability to leverage clade-specific mutations across the entire RSV genome, whereas our H3N2 tool, as well as Freyja, only utilizes the HA segment to call variants. Thus, a key objective for future surveillance efforts is the development of a robust IAV deconvolution tool capable of leveraging clade-defining mutations distributed across all IAV H3N2 genome segments. This strategy would facilitate accurate clade assignment even in instances of low or absent coverage of the hemagglutinin (HA) gene, by incorporating nucleotide polymorphisms from other genomic regions. Such an approach would enhance the reliability and resolution of lineage determination in metagenomic datasets derived from complex environmental samples, such as wastewater.

Additionally, the scope and effectiveness of an influenza and respiratory virus surveillance program would be strengthened by incorporating a comprehensive, full-genome sequencing assay capable of detecting H1N1(pdm09) viruses. Given that H1N1(pdm09) co-circulates with H3N2 and RSV B co-circulates with RSV A, simultaneous detection and characterization of these pathogens is crucial for a more complete understanding of community-level viral dynamics. In the present study, the exclusion of H1N1(pdm09) and RSV B detection was a limitation due to resource constraints. Future work should prioritize inclusion of these targets to address current gaps in our WBS methodology. Development of an H1N1(pdm09) targeted assay is of particular importance as this is a previous pandemic swine flu strain which poses a potential future threat, as mutations which arise in this strain could result in increased transmissibility and worsened disease outcomes.

## Conclusion

In summary, this study demonstrates the feasibility and efficacy of wastewater-based genomic surveillance for monitoring the circulation and diversity of respiratory viruses, specifically IAV H3N2 and RSV A. Through comparative analysis of the variant-calling tools Alvoc and Freyja, we show that both are capable of deconvoluting mixed viral populations in environmental samples, with Alvoc exhibiting improved fine-resolution of RSV A lineage assignments. These findings highlight the potential of wastewater genomics to complement traditional clinical surveillance by providing real-time, population-level data on variant prevalence and abundance. To enhance the scope and utility of future surveillance efforts, comprehensive deconvolution tools that utilize clade-defining mutations from all Influenza A genome segments, along with expanded assay panels to include additional co-circulating H1N1(pdm09) and RSV B genomes should be developed. Furthermore, optimization of virus concentration, isolation, and sequencing workflows will be essential to improve genome recovery and assay sensitivity. Together, these advancements will strengthen the capacity of wastewater-based surveillance systems to inform public health decision-making, guide IAV vaccine strain selection, predict RSV A vaccine escape variants from caused by mutations, and support the early detection of emerging variants with public health significance.

## Acknowledgments

The authors of this manuscript would like to thank and acknowledge the support of The Regional Municipality of York, Region of Waterloo Public Health and Paramedic Services, Regional Municipality of Peel, and Hamilton Water in the evaluation of these assays through providing samples for testing.

## Data Availability

Raw data from all samples are available on the SRA database under Bioproject accession number PRJNA1288507.

## References

1. Del Riccio M, Caini S, Bonaccorsi G, Lorini C, Paget J, Van Der Velden K, et al. Global analysis of respiratory viral circulation and timing of epidemics in the pre–COVID-19 and COVID-19 pandemic eras, based on data from the Global Influenza Surveillance and Response System (GISRS). Int J Infect Dis. 2024 Jul;144:107052.

2. Bloom-Feshbach K, Alonso WJ, Charu V, Tamerius J, Simonsen L, Miller MA, et al. Latitudinal Variations in Seasonal Activity of Influenza and Respiratory Syncytial Virus (RSV): A Global Comparative Review. Cowling BJ, editor. PLoS ONE. 2013 Feb 14;8(2):e54445.

3. Habbous S, Hota S, Allen VG, Henry M, Hellsten E. Changes in hospitalizations and emergency department respiratory viral diagnosis trends before and during the COVID-19 pandemic in Ontario, Canada. Wu AG, editor. PLOS ONE. 2023 Jun 16;18(6):e0287395.

4. Zimmerman RK, Balasubramani GK, D’Agostino HEA, Clarke L, Yassin M, Middleton DB, et al. Population-based hospitalization burden estimates for respiratory viruses, 2015– 2019. Influenza Other Respir Viruses. 2022 Nov;16(6):1133–40.

5. Griffiths C, Drews SJ, Marchant DJ. Respiratory Syncytial Virus: Infection, Detection, and New Options for Prevention and Treatment. Clin Microbiol Rev. 2017 Jan;30(1):277–319.

6. Javanian M, Barary M, Ghebrehewet S, Koppolu V, Vasigala V, Ebrahimpour S. A brief review of influenza virus infection. J Med Virol. 2021 Aug;93(8):4638–46.

7. Ambrosch A, Luber D, Klawonn F, Kabesch M. Focusing on severe infections with the respiratory syncytial virus (RSV) in adults: Risk factors, symptomatology and clinical course compared to influenza A / B and the original SARS-CoV-2 strain. J Clin Virol. 2023 Apr;161:105399.

8. Influenza (seasonal) [Internet]. World Helath Organization; 2025. Available from: https://www.who.int/news-room/fact-sheets/detail/influenza-(seasonal)

9. Li Y, Wang X, Blau DM, Caballero MT, Feikin DR, Gill CJ, et al. Global, regional, and national disease burden estimates of acute lower respiratory infections due to respiratory syncytial virus in children younger than 5 years in 2019: a systematic analysis. The Lancet. 2022 May;399(10340):2047–64.

10. Respiratory syncytial virus (RSV) [Internet]. World Helath Organization; 2025. Available from: https://www.who.int/news-room/fact-sheets/detail/respiratory-syncytial-virus-%28rsv%29

11. Macias AE, McElhaney JE, Chaves SS, Nealon J, Nunes MC, Samson SI, et al. The disease burden of influenza beyond respiratory illness. Vaccine. 2021 Mar;39:A6–14.

12. Averin A, Atwood M, Sato R, Yacisin K, Begier E, Shea K, et al. Attributable Cost of Adult Respiratory Syncytial Virus Illness Beyond the Acute Phase. Open Forum Infect Dis. 2024 Feb 29;11(3):ofae097.

13. McCarthy Z, Xu S, Rahman A, Bragazzi NL, Corrales-Medina VF, Lee J, et al. Modelling the linkage between influenza infection and cardiovascular events via thrombosis. Sci Rep. 2020 Aug 31;10(1):14264.

14. Xie Y, Choi T, Al-Aly Z. Long-term outcomes following hospital admission for COVID-19 versus seasonal influenza: a cohort study. Lancet Infect Dis. 2024 Mar;24(3):239–55.

15. Blanken MO, Rovers MM, Molenaar JM, Winkler-Seinstra PL, Meijer A, Kimpen JLL, et al. Respiratory Syncytial Virus and Recurrent Wheeze in Healthy Preterm Infants. N Engl J Med. 2013 May 9;368(19):1791–9.

16. Trautmannsberger I, Plagg B, Adamek I, Mader S, De Luca D, Esposito S, et al. The Multifaceted Burden of Respiratory Syncytial Virus (RSV) Infections in Young Children on the Family: A European Study. Infect Dis Ther [Internet]. 2024 May 20 [cited 2024 Jun 11]; Available from: https://link.springer.com/10.1007/s40121-024-00989-0

17. Blanchet Zumofen MH, Frimpter J, Hansen SA. Impact of Influenza and Influenza-Like Illness on Work Productivity Outcomes: A Systematic Literature Review. PharmacoEconomics. 2023 Mar;41(3):253–73.

18. Cohen C, Zar HJ. Early life respiratory syncytial virus disease—a preventable burden. Lancet Infect Dis. 2024 Apr;S1473309924002615.

19. Tran PT, Nduaguba SO, Wang Y, Diaby V, Finelli L, Choi Y, et al. Economic Burden of Medically Attended Respiratory Syncytial Virus Infections Among Privately Insured Children Under 5 Years of Age in the USA. Influenza Other Respir Viruses. 2024 Jul;18(7):e13347.

20. Yan S, Weycker D, Sokolowski S. US healthcare costs attributable to type A and type B influenza. Hum Vaccines Immunother. 2017 Sep 2;13(9):2041–7.

21. Herring WL, Zhang Y, Shinde V, Stoddard J, Talbird SE, Rosen B. Clinical and economic outcomes associated with respiratory syncytial virus vaccination in older adults in the United States. Vaccine. 2022 Jan;40(3):483–93.

22. Belongia EA, McLean HQ. Influenza Vaccine Effectiveness: Defining the H3N2 Problem. Clin Infect Dis. 2019 Oct 30;69(10):1817–23.

23. Schmidt K, Ben Moussa M, Buckrell S, Rahal A, Chestley T, Bastien N, et al. National Influenza Annual Report, Canada, 2022–2023: Canada’s first fall epidemic since the 2019– 2020 season. Can Commun Dis Rep. 2023 Nov 20;49(10):413–24.

24. Ben Moussa M, Nwosu A, Schmidt K, Buckrell S, Rahal A, Lee L, et al. National Influenza Annual Report 2023–2024: A focus on influenza B and public health implications. Can Commun Dis Rep. 2024 Nov 7;50(11):393–9.

25. Bøås H, Havdal LB, Størdal K, Døllner H, Leegaard TM, Bekkevold T, et al. No association between disease severity and respiratory syncytial virus subtypes RSV-A and RSV-B in hospitalized young children in Norway. Miron VD, editor. PLOS ONE. 2024 Mar 11;19(3):e0298104.

26. Ciarlitto C, Vittucci AC, Antilici L, Concato C, Di Camillo C, Zangari P, et al. Respiratory Syncityal Virus A and B: three bronchiolitis seasons in a third level hospital in Italy. Ital J Pediatr. 2019 Dec;45(1):115.

27. Chen J, Qiu X, Avadhanula V, Shepard SS, Kim D, Hixson J, et al. Novel and extendable genotyping system for human respiratory syncytial virus based on whole-genome sequence analysis. Influenza Other Respir Viruses. 2022 May;16(3):492–500.

28. Goya S, Galiano M, Nauwelaers I, Trento A, Openshaw PJ, Mistchenko AS, et al. Toward unified molecular surveillance of RSV: A proposal for genotype definition. Influenza Other Respir Viruses. 2020 May;14(3):274–85.

29. Goya S, Ruis C, Neher RA, Meijer A, Aziz A, Hinrichs AS, et al. Standardized Phylogenetic Classification of Human Respiratory Syncytial Virus Below the Subgroup Level. Emerg Infect Dis. 2024 Aug;30(8).

30. Ladner JT, Sahl JW. Towards a post-pandemic future for global pathogen genome sequencing. PLOS Biol. 2023 Aug 1;21(8):e3002225.

31. Williams TGS, Snell LB, Alder C, Charalampous T, Alcolea-Medina A, Sehmi JK, et al. Feasibility and clinical utility of local rapid Nanopore influenza A virus whole genome sequencing for integrated outbreak management, genotypic resistance detection and timely surveillance. Microb Genomics. 2023 Aug 17;9(8).

32. Galli C, Ebranati E, Pellegrinelli L, Airoldi M, Veo C, Della Ventura C, et al. From Clinical Specimen to Whole Genome Sequencing of A(H3N2) Influenza Viruses: A Fast and Reliable High-Throughput Protocol. Vaccines. 2022 Aug 19;10(8):1359.

33. MacFadden DR, McGeer A, Athey T, Perusini S, Olsha R, Li A, et al. Use of Genome Sequencing to Define Institutional Influenza Outbreaks, Toronto, Ontario, Canada, 2014–15. Emerg Infect Dis. 2018 Mar;24(3):492–7.

34. Vemula S, Zhao J, Liu J, Wang X, Biswas S, Hewlett I. Current Approaches for Diagnosis of Influenza Virus Infections in Humans. Viruses. 2016 Apr 12;8(4):96.

35. Fall A, Han L, Yunker M, Gong YN, Li TJ, Norton JM, et al. Evolution of Influenza A(H3N2) Viruses in 2 Consecutive Seasons of Genomic Surveillance, 2021–2023. Open Forum Infect Dis. 2023 Dec 1;10(12):ofad577.

36. May F, Ginige S, Firman E, Li YS, Soonarane YK, Smoll N, et al. Estimating the incidence of COVID-19, influenza and respiratory syncytial virus infection in three regions of Queensland, Australia, winter 2022: findings from a novel longitudinal testing-based sentinel surveillance programme. BMJ Open. 2024 Apr;14(4):e081793.

37. Toribio-Avedillo D, Gómez-Gómez C, Sala-Comorera L, Rodríguez-Rubio L, Carcereny A, García-Pedemonte D, et al. Monitoring influenza and respiratory syncytial virus in wastewater. Beyond COVID-19. Sci Total Environ. 2023 Sep;892:164495.

38. Parkins MD, Lee BE, Acosta N, Bautista M, Hubert CRJ, Hrudey SE, et al. Wastewater-based surveillance as a tool for public health action: SARS-CoV-2 and beyond. Forrest GN, editor. Clin Microbiol Rev. 2024 Mar 14;37(1):e00103–22.

39. Hayes EK, Gouthro MT, LeBlanc JJ, Gagnon GA. Simultaneous detection of SARS-CoV-2, influenza A, respiratory syncytial virus, and measles in wastewater by multiplex RT-qPCR. Sci Total Environ. 2023 Sep;889:164261.

40. Lee AJ, Carson S, Reyne MI, Marshall A, Moody D, Allen DM, et al. Wastewater monitoring of human and avian influenza A viruses in Northern Ireland: a genomic surveillance study. Lancet Microbe. 2024 Dec;5(12):100933.

41. Kitajima M, Ahmed W, Bibby K, Carducci A, Gerba CP, Hamilton KA, et al. SARS-CoV-2 in wastewater: State of the knowledge and research needs. Sci Total Environ. 2020 Oct;739:139076.

42. Keshaviah A, Diamond MB, Wade MJ, Scarpino SV, Ahmed W, Amman F, et al. Wastewater monitoring can anchor global disease surveillance systems. Lancet Glob Health. 2023 Jun;11(6):e976–81.

43. Berry I, Brown KA, Buchan SA, Hohenadel K, Kwong JC, Patel S, et al. A better normal in Canada will need a better detection system for emerging and re-emerging respiratory pathogens. Can Med Assoc J. 2022 Sep 19;194(36):E1250–4.

44. Thampi N, Mercier E, Paes B, Edwards JO, Rodgers-Gray B, Delatolla R. Perspective: the potential of wastewater-based surveillance as an economically feasible game changer in reducing the global burden of pediatric respiratory syncytial virus infection. Front Public Health. 2024 Jan 12;11:1316531.

45. Mercier E, Pisharody L, Guy F, Wan S, Hegazy N, D’Aoust PM, et al. Wastewater-based surveillance identifies start to the pediatric respiratory syncytial virus season in two cities in Ontario, Canada. Front Public Health. 2023 Sep 26;11:1261165.

46. Zulli A, Varkila MRJ, Parsonnet J, Wolfe MK, Boehm AB. Observations of Respiratory Syncytial Virus (RSV) Nucleic Acids in Wastewater Solids Across the United States in the 2022–2023 Season: Relationships with RSV Infection Positivity and Hospitalization Rates. ACS EST Water. 2024 Apr 12;4(4):1657–67.

47. Nakauchi M, Yasui Y, Miyoshi T, Minagawa H, Tanaka T, Tashiro M, et al. One-step real-time reverse transcription-PCR assays for detecting and subtyping pandemic influenza A/H1N1 2009, seasonal influenza A/H1N1, and seasonal influenza A/H3N2 viruses. J Virol Methods. 2011 Jan;171(1):156–62.

48. Zafeiriadou A, Kaltsis L, Thomaidis NS, Markou A. Simultaneous detection of influenza A, B and respiratory syncytial virus in wastewater samples by one-step multiplex RT-ddPCR assay. Hum Genomics. 2024 May 20;18(1):48.

49. Malla B, Shrestha S, Haramoto E. Optimization of the 5-plex digital PCR workflow for simultaneous monitoring of SARS-CoV-2 and other pathogenic viruses in wastewater. Sci Total Environ. 2024 Feb;913:169746.

50. Allen DM, Reyne MI, Allingham P, Levickas A, Bell SH, Lock J, et al. Genomic Analysis and Surveillance of Respiratory Syncytial Virus Using Wastewater-Based Epidemiology. J Infect Dis. 2024 Apr 18;jiae205.

51. Quick J, Grubaugh ND, Pullan ST, Claro IM, Smith AD, Gangavarapu K, et al. Multiplex PCR method for MinION and Illumina sequencing of Zika and other virus genomes directly from clinical samples. Nat Protoc. 2017 Jun;12(6):1261–6.

52. Quick J, Lansdowne L. ARTIC SARS-CoV-2 sequencing protocol v4 (LSK114) V.4. Protocols.io; 2024.

53. Tyson JR, James P, Stoddart D, Sparks N, Wickenhagen A, Hall G, et al. Improvements to the ARTIC multiplex PCR method for SARS-CoV-2 genome sequencing using nanopore. Genomics; 2020.

54. Williams RC, Farkas K, Garcia-Delgado A, Adwan L, Kevill JL, Cross G, et al. Simultaneous detection and characterization of common respiratory pathogens in wastewater through genomic sequencing. Water Res. 2024 Jun;256:121612.

55. Vo V, Harrington A, Chang CL, Baker H, Moshi MA, Ghani N, et al. Identification and genome sequencing of an influenza H3N2 variant in wastewater from elementary schools during a surge of influenza A cases in Las Vegas, Nevada. Sci Total Environ. 2023 May;872:162058.

56. Ladner JT, Sahl JW. Towards a post-pandemic future for global pathogen genome sequencing. PLOS Biol. 2023 Aug 1;21(8):e3002225.

57. Toribio-Avedillo D, Gómez-Gómez C, Sala-Comorera L, Galofré B, Muniesa M. Adapted methods for monitoring influenza virus and respiratory syncytial virus in sludge and wastewater. Sci Total Environ. 2024 Mar;918:170636.

58. Hughes B, Duong D, White BJ, Wigginton KR, Chan EMG, Wolfe MK, et al. Respiratory Syncytial Virus (RSV) RNA in Wastewater Settled Solids Reflects RSV Clinical Positivity Rates. Environ Sci Technol Lett. 2022 Feb 8;9(2):173–8.

59. Rector A, Bloemen M, Thijssen M, Pussig B, Beuselinck K, Van Ranst M, et al. Respiratory Viruses in Wastewater Compared with Clinical Samples, Leuven, Belgium. Emerg Infect Dis. 2024 Jan;30(1).

60. Li H, He F, Lv Z, Yi L, Zhang Z, Li H, et al. Tailored wastewater surveillance framework uncovered the epidemics of key pathogens in a Northwestern city of China. Sci Total Environ. 2024 May;926:171833.

61. Boehm AB, Hughes B, Duong D, Chan-Herur V, Buchman A, Wolfe MK, et al. Wastewater concentrations of human influenza, metapneumovirus, parainfluenza, respiratory syncytial virus, rhinovirus, and seasonal coronavirus nucleic-acids during the COVID-19 pandemic: a surveillance study. Lancet Microbe. 2023 May;4(5):e340–8.

62. De Melo T, Islam G, Simmons DBD, Desaulniers JP, Kirkwood AE. An alternative method for monitoring and interpreting influenza A in communities using wastewater surveillance. Front Public Health. 2023 Jul 27;11:1141136.

63. DeJonge PM, Adams C, Pray I, Schussman MK, Fahney RB, Shafer M, et al. Wastewater Surveillance Data as a Complement to Emergency Department Visit Data for Tracking Incidence of Influenza A and Respiratory Syncytial Virus — Wisconsin, August 2022– March 2023. MMWR Morb Mortal Wkly Rep. 2023 Sep 15;72(37):1005–9.

64. Faherty EAG, Yuce D, Korban C, Bemis K, Kowalski R, Gretsch S, et al. Correlation of wastewater surveillance data with traditional influenza surveillance measures in Cook County, Illinois, October 2022–April 2023. Sci Total Environ. 2024 Feb;912:169551.

65. Germano ER, Flores T, Freed GS, Kim K, Tulinsky GH, Yang A, et al. Building-level wastewater surveillance localizes interseasonal influenza variation. Imperiale MJ, editor. mSphere. 2024 Jan 30;9(1):e00600–23.

66. Girón-Guzmán I, Cuevas-Ferrando E, Barranquero R, Díaz-Reolid A, Puchades-Colera P, Falcó I, et al. Urban wastewater-based epidemiology for multi-viral pathogen surveillance in the Valencian region, Spain. Water Res. 2024 May;255:121463.

67. Dlamini M, Msolo L, Ehi Ebomah K, Nontongana N, Ifeanyi Okoh A. A systematic review on the incidence of influenza viruses in wastewater matrices: Implications for public health. Hemati S, editor. PLOS ONE. 2024 Apr 25;19(4):e0291900.

68. Ando H, Ahmed W, Iwamoto R, Ando Y, Okabe S, Kitajima M. Impact of the COVID-19 pandemic on the prevalence of influenza A and respiratory syncytial viruses elucidated by wastewater-based epidemiology. Sci Total Environ. 2023 Jul;880:162694.

69. Boehm AB, Wolfe MK, White BJ, Hughes B, Duong D, Bidwell A. More than a Tripledemic: Influenza A Virus, Respiratory Syncytial Virus, SARS-CoV-2, and Human Metapneumovirus in Wastewater during Winter 2022–2023. Environ Sci Technol Lett. 2023 Aug 8;10(8):622–7.

70. Lehto KM, Länsivaara A, Hyder R, Luomala O, Lipponen A, Hokajärvi AM, et al. Wastewater-based surveillance is an efficient monitoring tool for tracking influenza A in the community. Water Res. 2024 Jun;257:121650.

71. Markt R, Stillebacher F, Nägele F, Kammerer A, Peer N, Payr M, et al. Expanding the Pathogen Panel in Wastewater Epidemiology to Influenza and Norovirus. Viruses. 2023 Jan 17;15(2):263.

72. Mercier E, D’Aoust PM, Thakali O, Hegazy N, Jia JJ, Zhang Z, et al. Municipal and neighbourhood level wastewater surveillance and subtyping of an influenza virus outbreak. Sci Rep. 2022 Sep 22;12(1):15777.

73. Raya S, Malla B, Shrestha S, Sthapit N, Kattel H, Sharma ST, et al. Quantification of multiple respiratory viruses in wastewater in the Kathmandu Valley, Nepal: Potential implications of wastewater-based epidemiology for community disease surveillance in developing countries. Sci Total Environ. 2024 Apr;920:170845.

74. Wolken M, Sun T, McCall C, Schneider R, Caton K, Hundley C, et al. Wastewater surveillance of SARS-CoV-2 and influenza in preK-12 schools shows school, community, and citywide infections. Water Res. 2023 Mar;231:119648.

75. Zafeiriadou A, Kaltsis L, Kostakis M, Kapes V, Thomaidis NS, Markou A. Wastewater surveillance of the most common circulating respiratory viruses in Athens: The impact of COVID-19 on their seasonality. Sci Total Environ. 2023 Nov;900:166136.

76. Dumke R, Geissler M, Skupin A, Helm B, Mayer R, Schubert S, et al. Simultaneous Detection of SARS-CoV-2 and Influenza Virus in Wastewater of Two Cities in Southeastern Germany, January to May 2022. Int J Environ Res Public Health. 2022 Oct 17;19(20):13374.

77. Heijnen L, Medema G. Surveillance of Influenza A and the pandemic influenza A (H1N1) 2009 in sewage and surface water in the Netherlands. J Water Health. 2011 Sep 1;9(3):434–42.

78. Stockdale SR, Blanchard AM, Nayak A, Husain A, Nashine R, Dudani H, et al. RNA-Seq of untreated wastewater to assess COVID-19 and emerging and endemic viruses for public health surveillance. Lancet Reg Health - Southeast Asia. 2023 Jul;14:100205.

79. Tisza M, Javornik Cregeen S, Avadhanula V, Zhang P, Ayvaz T, Feliz K, et al. Wastewater sequencing reveals community and variant dynamics of the collective human virome. Nat Commun. 2023 Oct 28;14(1):6878.

80. Wolfe MK, Duong D, Bakker KM, Ammerman M, Mortenson L, Hughes B, et al. Wastewater-Based Detection of Two Influenza Outbreaks. Environ Sci Technol Lett. 2022 Aug 9;9(8):687–92.

81. Freyja: Depth-Weighted De-Mixing [Internet]. Github; Available from: https://github.com/andersen-lab/Freyja

82. Ferdous J, Kunkleman S, Taylor W, Harris A, Gibas CJ, Schlueter JA. A gold standard dataset and evaluation of methods for lineage abundance estimation from wastewater. Sci Total Environ. 2024 Oct;948:174515.

83. Sutcliffe SG, Kraemer SA, Ellmen I, Knapp JJ, Overton AK, Nash D, et al. Tracking SARS-CoV-2 variants of concern in wastewater: an assessment of nine computational tools using simulated genomic data. Microb Genomics. 2024 May 24;10(5).

84. Karthikeyan S, Levy JI, De Hoff P, Humphrey G, Birmingham A, Jepsen K, et al. Wastewater sequencing reveals early cryptic SARS-CoV-2 variant transmission. Nature. 2022 Sep 1;609(7925):101–8.

85. Alvoc: Abundance learning for variants of concern [Internet]. Github; Available from: https://github.com/alvoc/alvoc

86. Ellmen I, Lynch MDJ, Nash D, Cheng J, Nissimov JI, Charles TC. Alcov: Estimating Variant of Concern Abundance from SARS-CoV-2 Wastewater Sequencing Data. Health Informatics; 2021.

87. Zhou B, Donnelly ME, Scholes DT, St. George K, Hatta M, Kawaoka Y, et al. Single-Reaction Genomic Amplification Accelerates Sequencing and Vaccine Production for Classical and Swine Origin Human Influenza A Viruses. J Virol. 2009 Oct;83(19):10309– 13.

88. Dong X, Deng YM, Aziz A, Whitney P, Clark J, Harris P, et al. A simplified, amplicon-based method for whole genome sequencing of human respiratory syncytial viruses. J Clin Virol. 2023 Apr;161:105423.

89. Köndgen S, Oh DY, Thürmer A, Sedaghatjoo S, Patrono LV, Calvignac-Spencer S, et al. A robust, scalable, and cost-efficient approach to whole genome sequencing of RSV directly from clinical samples. Dekker JP, editor. J Clin Microbiol. 2024 Mar 13;62(3):e01111–23.

90. Croville G, Walch M, Sécula A, Lèbre L, Silva S, Filaire F, et al. An amplicon-based nanopore sequencing workflow for rapid tracking of avian influenza outbreaks, France, 2020-2022. Front Cell Infect Microbiol. 2024 Jan 22;14:1257586.

91. Diaz A, Enomoto S, Romagosa A, Sreevatsan S, Nelson M, Culhane M, et al. Genome plasticity of triple-reassortant H1N1 influenza A virus during infection of vaccinated pigs. J Gen Virol. 2015 Oct 1;96(10):2982–93.

92. Cruz CD, Icochea ME, Espejo V, Troncos G, Castro-Sanguinetti GR, Schilling MA, et al. Highly Pathogenic Avian Influenza A(H5N1) from Wild Birds, Poultry, and Mammals, Peru. Emerg Infect Dis. 2023 Dec;29(12):2572–6.

93. Dadonaite B, Gilbertson B, Knight ML, Trifkovic S, Rockman S, Laederach A, et al. The structure of the influenza A virus genome. Nat Microbiol. 2019;4(11).

94. Al-leimon O, Shihadeh H, Yousef AA, Khraim A, Siwwad R. Respiratory syncytial virus: A review of current basic and clinical knowledge. Qatar Med J. 2024 Jan;2024(4).

95. Chen S. Ultrafast one-pass FASTQ data preprocessing, quality control, and deduplication using fastp. iMeta. 2023 May;2(2):e107.

96. Chen S, Zhou Y, Chen Y, Gu J. fastp: an ultra-fast all-in-one FASTQ preprocessor. Bioinformatics. 2018 Sep 1;34(17):i884–90.

97. Li H, Durbin R. Fast and accurate long-read alignment with Burrows–Wheeler transform. Bioinformatics. 2010 Mar 1;26(5):589–95.

98. Danecek P, Bonfield JK, Liddle J, Marshall J, Ohan V, Pollard MO, et al. Twelve years of SAMtools and BCFtools. GigaScience. 2021 Feb;10(2).

99. nh13/DWGSIM: Whole Genome Simulator for Next-Generation Sequencing [Internet]. Available from: https://github.com/nh13/DWGSIM/tree/main

100. Martin M. Cutadapt Removes Adapter Sequences From High-Throughput Sequencing Reads. EMBnet.journal. 2011;17(1):10–2.

101. Li H. New strategies to improve minimap2 alignment accuracy. Alkan C, editor. Bioinformatics. 2021 Dec 7;37(23):4572–4.

102. Zhang M, Roldan-Hernandez L, Boehm A. Persistence of human respiratory viral RNA in wastewater-settled solids. Elkins CA, editor. Appl Environ Microbiol. 2024 Apr 17;90(4):e02272–23.

103. Skowronski DM, Zhan Y, Kaweski SE, Sabaiduc S, Khalid A, Olsha R, et al. 2023/24 mid-season influenza and Omicron XBB.1.5 vaccine effectiveness estimates from the Canadian Sentinel Practitioner Surveillance Network (SPSN). Eurosurveillance. 2024 Feb 15;29(7).

104. Khare S, Gurry C, Freitas L, B Schultz M, Bach G, Diallo A, et al. GISAID’s Role in Pandemic Response. China CDC Wkly. 2021;3(49):1049–51.

105. Zhang J, Zhang L, Li J, Ma J, Wang Y, Sun Y, et al. Moderate effectiveness of influenza vaccine in outpatient settings: A test-negative study in Beijing, China, 2023/24 season. Vaccine. 2025 Feb;46:126662.

106. Martínez-Baz I, Navascués A, Trobajo-Sanmartín C, Pozo F, Fernández-Huerta M, Olazabal-Arruiz M, et al. Effectiveness of influenza vaccination in preventing confirmed influenza cases and hospitalizations in Northern Spain, 2023/24 season: A population-based test-negative case-control study. Int J Infect Dis. 2025 Feb;151:107364.

107. Marron L, McKenna A, O’Donnell J, Joyce M, Bennett C, Connell J, et al. Influenza Vaccine Effectiveness Against Symptomatic Influenza in Primary Care: A Test Negative Case Control Study Over Two Influenza Seasons 2022/2023 and 2023/2024 in Ireland. Influenza Other Respir Viruses. 2024 Dec;18(12):e70023.

108. Separovic L, Zhan Y, Kaweski SE, Sabaiduc S, Carazo S, Olsha R, et al. Interim estimates of vaccine effectiveness against influenza A(H1N1)pdm09 and A(H3N2) during a delayed influenza season, Canada, 2024/25. Eurosurveillance. 2025 Jan 30;30(4).

109. Centers of Disease Control and Protection. ACIP Evidence to Recommendations for Use of Protein Subunit RSV vaccines (GSK Arexvy or Pfizer Abrysvo) in All Adults Aged ≥75 years and in Adults Aged 60–74 at Increased Risk of Severe RSV Disease. U.S. Department of Health and Human Services.

110. Zhu Q, Patel NK, McAuliffe JM, Zhu W, Wachter L, McCarthy MP, et al. Natural Polymorphisms and Resistance-Associated Mutations in the Fusion Protein of Respiratory Syncytial Virus (RSV): Effects on RSV Susceptibility to Palivizumab. J Infect Dis. 2012 Feb 15;205(4):635–8.

111. Yasui Y, Yamaji Y, Sawada A, Ito T, Nakayama T. Cell fusion assay by expression of respiratory syncytial virus (RSV) fusion protein to analyze the mutation of palivizumab-resistant strains. J Virol Methods. 2016 May;231:48–55.

112. Holland LA, Holland SC, Smith MF, Leonard VR, Murugan V, Nordstrom L, et al. Genomic Sequencing Surveillance to Identify Respiratory Syncytial Virus Mutations, Arizona, USA. Emerg Infect Dis. 2023 Nov;29(11).

113. Farkas K, Pellett C, Alex-Sanders N, Bridgman MTP, Corbishley A, Grimsley JMS, et al. Comparative Assessment of Filtration- and Precipitation-Based Methods for the Concentration of SARS-CoV-2 and Other Viruses from Wastewater. Faucher SP, editor. Microbiol Spectr. 2022 Aug 31;10(4):e01102–22.

114. Shu B, Kirby MK, Davis WG, Warnes C, Liddell J, Liu J, et al. Multiplex Real-Time Reverse Transcription PCR for Influenza A Virus, Influenza B Virus, and Severe Acute Respiratory Syndrome Coronavirus 2. Emerg Infect Dis. 2021;27(7):1821–30.

115. US Center for Disease Control and Prevention. CDC’s influenza SARS-CoV-2 multiplex assay. US Center for Disease Control and Prevention. Available from: https://www.cdc.gov/flu/php/laboratories/influenza-sars-cov-2-multiplex-assay.html

