## Supplementary Tables and Figure for "A Genome Sequence Variant Monitoring Program for Seasonal Influenza A H3N2 and Respiratory Syncytial Virus A using Wastewater-Based Surveillance in Ontario, Canada"

### *Supplementary Material*

**Supplementary Table 1**. *Locations of Wastewater Sample Collection*.

| Region | Collection Site | Collection System | Sample Type |
|---|---|---|---|
| Peel | GE Booth | Wastewater Treatment Plant | 24 h composite |
| York | Leslie Street (York 1) | Linear Pipe Collection Site | Grab |
| York | Warden (York 2) | Pumping Station | Grab |
| Waterloo | Kitchener | Wastewater Treatment Plant | 24 h composite |
| Waterloo | Cambridge | Wastewater Treatment Plant | 24 h composite |
| Hamilton | Hamilton | Wastewater Treatment Plant | 24 h composite |

**Supplementary Table 2.** *Dates of Wastewater Sample Collection.* The collection date of samples from six sites and dates of surveillance week.

| Surveillance Week | Dates of Surveillance Week | GE Booth | York 1 | York 2 | Kitchener | Cambridge | Hamilton |
|---|---|---|---|---|---|---|---|
| 45 | Nov 5 - Nov 11, 2023 | Nov 6, 2023 | Nov 6, 2023 | Nov 6, 2023 | Nov 6, 2023 | Nov 6, 2023 | Nov 7, 2023 |
| 46 | Nov 12 - Nov 18, 2023 | Nov 13, 2023 | Nov 13, 2023 | Nov 13, 2023 | - | - | Nov 14, 2023 |
| 47 | Nov 19 - Nov 25, 2023 | Nov 20, 2023 | Nov 20, 2023 | Nov 20, 2023 | Nov 20, 2023 | Nov 20, 2023 | Nov 21, 2023 |
| 48 | Nov 26 - Dec 2, 2023 | Nov 27, 2023 | Nov 27, 2023 | Nov 27, 2023 | Nov 27, 2023 | Nov 27, 2023 | Nov 28, 2023 |
| 49 | Dec 3 - Dec 9, 2023 | Dec 4, 2023 | Dec 4, 2023 | Dec 4, 2023 | Dec 4, 2023 | Dec 4, 2023 | Dec 5, 2023 |
| 50 | Dec 10 - Dec 16, 2023 | Dec 11, 2023 | Dec 11, 2023 | Dec 11, 2023 | Dec 11, 2023 | Dec 11, 2023 | Dec 12, 2023 |
| 51 | Dec 17 - Dec 23, 2023 | Dec 18, 2023 | Dec 18, 2023 | Dec 18, 2023 | Dec 18, 2023 | Dec 18, 2023 | Dec 15, 2023 |
| 52 | Dec 24 - Dec 30, 2023 | Dec 26, 2023 | - | - | - | Dec 25, 2023 | Dec 27, 2023 |
| 1 | Dec 31, 2023 - Jan 6, 2024 | Jan 3, 2024 | Jan 3, 2024 | Jan 3, 2024 | Jan 1, 2024 | - | Jan 2, 2024 |
| 2 | Jan 7 - Jan 13, 2024 | Jan 8, 2024 | Jan 8, 2024 | Jan 8, 2024 | Jan 15, 2024 | - | Jan 9, 2024 |
| 3 | Jan 14 - Jan 20, 2024 | Jan 17, 2024 | Jan 15, 2024 | Jan 15, 2024 | Jan 17, 2024 | Jan 15, 2024 | Jan 16, 2024 |
| 4 | Jan 21 - Jan 27, 2024 | Jan 22, 2024 | Jan 22, 2024 | Jan 22, 2024 | Jan 22, 2024 | Jan 22, 2024 | Jan 23, 2024 |
| 5 | Jan 28 - Feb 3, 2024 | Jan 29, 2024 | Jan 29, 2024 | Jan 29, 2024 | Jan 29, 2024 | Jan 29, 2024 | Jan 30, 2024 |

**Supplementary Table 3.** *IAV H3N2 Primers*. List of tiled-amplicon IAV H3N2 primers, including, name, amplicon, sequence, size, %GC, melting temperature, segment name, strand, primer pool, and pool concentration.

| Name | Amplicon | Seq | size | %gc | Tm | Segment Name | Strand | Pool | [pool] (uM) |
|---|---|---|---|---|---|---|---|---|---|
| PB2_1_LEFT | 1 | AAACCACAGTGGACCATATGGC | 22 | 50 | 61.07 | PB2 | + | 1 | 0.4 |
| PB2_1_RIGHT | 1 | TTGGCACTGAGGTCTGCATGA | 21 | 52.38 | 62.1 | PB2 | - | 1 | 0.4 |
| PB2_2_LEFT | 2 | AACCTTTGGCCCTGTCCATTTT | 22 | 45.45 | 61.15 | PB2 | + | 2 | 0.2 |
| PB2_2_RIGHT | 2 | GGCCGCAATAATTAGGCTTTGG | 22 | 50 | 60.72 | PB2 | - | 2 | 0.2 |
| PB2_3_LEFT | 3 | AGCCAGAATACTAACATCAGAATCACA | 27 | 37.04 | 60.86 | PB2 | + | 1 | 0.2 |
| PB2_3_RIGHT | 3 | AGCTTGTTCTTCAGTCGGGTTC | 22 | 50 | 60.99 | PB2 | - | 1 | 0.2 |
| PB2_4_LEFT | 4 | GCAGTGTCAGCAGATCCACT | 20 | 55 | 60.05 | PB2 | + | 2 | 0.4 |
| PB2_4_RIGHT | 4 | CTTGTGAAAACACCATGGCCAC | 22 | 50 | 60.98 | PB2 | - | 2 | 0.4 |
| PB2_5_LEFT | 5 | ACTAAGAAAAGCAACCAGAAGACTG | 25 | 40 | 59.85 | PB2 | + | 1 | 0.4 |
| PB2_5_RIGHT | 5 | TCTCGAACTCTCAAAAACCGATCA | 24 | 41.67 | 60.58 | PB2 | - | 1 | 0.4 |
| PB2_6_LEFT | 6 | AGGAATAAGAGTCAGCAAAATGGGT | 25 | 40 | 60.91 | PB2 | + | 2 | 0.4 |
| PB2_6_RIGHT | 6 | CCAAGTACGTCTCTCATTTGTTGG | 24 | 45.83 | 60.16 | PB2 | - | 2 | 0.4 |
| PB2_7_LEFT | 7 | AATCTTTAGTCCCCAAGGCCAC | 22 | 50 | 60.48 | PB2 | + | 1 | 0.2 |
| PB2_7_RIGHT | 7 | TTGATGCTTAATGCTGGTCCGT | 22 | 45.45 | 60.8 | PB2 | - | 1 | 0.2 |
| PB2_8_LEFT | 8 | ACTGACTGTGAATGTGAGGGGA | 22 | 50 | 61.21 | PB2 | + | 2 | 0.4 |
| PB2_8_RIGHT | 8 | GAAACAAGGTCGTTTTTAAACTATTCAGT | 29 | 31.03 | 60.17 | PB2 | - | 2 | 0.4 |
| PB1_1_LEFT | 9 | GTTCCAGCGCAAAATGCCATAA | 22 | 45.45 | 60.85 | PB1 | + | 1 | 0.4 |
| PB1_1_RIGHT | 9 | GACTTCTATGGTGTTGGCTAATGC | 24 | 45.83 | 60.28 | PB1 | - | 1 | 0.4 |
| PB1_2_LEFT | 10 | ATGGAAGCCGTTCAACAGACAA | 22 | 45.45 | 60.93 | PB1 | + | 2 | 0.4 |
| PB1_2_RIGHT | 10 | GGGTGTTGCAATAGCCCTTCTT | 22 | 50 | 61.33 | PB1 | - | 2 | 0.4 |
| PB1_3_LEFT | 11 | TGAATAAGAGAGGCTACCTAATAAGAGC | 28 | 39.29 | 60.61 | PB1 | + | 1 | 0.4 |
| PB1_3_RIGHT | 11 | ACATTATTGGTGCGATGCTCAG | 22 | 45.45 | 59.56 | PB1 | - | 1 | 0.4 |
| PB1_4_LEFT | 12 | AAAACCCCCGAATGTTTTTGGC | 22 | 45.45 | 60.93 | PB1 | + | 2 | 0.2 |
| PB1_4_RIGHT | 12 | ATGAGGGCAAAATCGTCTGAGG | 22 | 50 | 60.86 | PB1 | - | 2 | 0.2 |
| PB1_5_LEFT | 13 | ACATGCTAAGTACAGTTTTAGGAGTCT | 27 | 37.04 | 60.54 | PB1 | + | 1 | 0.3 |
| PB1_5_RIGHT | 13 | ATTTGGGCTGTTGCTGGTCC | 20 | 55 | 61.2 | PB1 | - | 1 | 0.3 |
| PB1_6_LEFT | 14 | GTCAGCTGACATGAGCATTGGA | 22 | 50 | 61.12 | PB1 | + | 2 | 0.2 |
| PB1_6_RIGHT | 14 | CCATGGGCTGGCATTACTACAG | 22 | 54.55 | 61.25 | PB1 | - | 2 | 0.2 |
| PB1_7_LEFT | 15 | CTTCACATCCCTGAAGTCTGCT | 22 | 50 | 60.21 | PB1 | + | 1 | 0.4 |
| PB1_7_RIGHT | 15 | AATCCTTCCAGACTCGAAGTCAA | 23 | 43.48 | 59.68 | PB1 | - | 1 | 0.4 |
| PA_1_LEFT | 16 | TGGAAGATTTTGTGCGACAATGC | 23 | 43.48 | 61.11 | PA | + | 1 | 0.4 |
| PA_1_RIGHT | 16 | CAAGGTAATATATGTGGACTTCTCTCCT | 28 | 39.29 | 60.56 | PA | - | 1 | 0.4 |
| PA_2_LEFT | 17 | GCAACACTACTGGAGCTGGAAA | 22 | 50 | 60.99 | PA | + | 2 | 0.4 |
| PA_2_RIGHT | 17 | GGCTCTAAAATTCTCAAGGCAGG | 23 | 47.83 | 59.87 | PA | - | 2 | 0.4 |
| PA_3_LEFT | 18 | AGAAAAAATTTGAAATCACAGGAACTATGC | 29 | 31.03 | 60.47 | PA | + | 1 | 0.4 |
| PA_3_RIGHT | 18 | TCCATGACAGCAGGTAATTTGAATT | 25 | 36 | 59.67 | PA | - | 1 | 0.4 |
| PA_4_LEFT | 19 | AGTGCATAAAAACATTCTTTGGATGGA | 27 | 33.33 | 60.75 | PA | + | 2 | 0.4 |
| PA_4_RIGHT | 19 | TTGCAATGTGTTCAATTGGGGC | 22 | 45.45 | 60.99 | PA | - | 2 | 0.4 |
| PA_5_LEFT | 20 | AACAAGGCCTGTGAGCTAACTG | 22 | 50 | 60.99 | PA | + | 1 | 0.4 |
| PA_5_RIGHT | 20 | TCTCAAGGACACAATATTTCTCCCA | 25 | 40 | 60.14 | PA | - | 1 | 0.4 |
| PA_6_LEFT | 21 | GTGGTAAACTTTGTGAGCATGGA | 23 | 43.48 | 59.68 | PA | + | 2 | 0.4 |
| PA_6_RIGHT | 21 | TGCATACAGGCTATTGAATACTGACT | 26 | 38.46 | 60.63 | PA | - | 2 | 0.4 |
| PA_7_LEFT | 22 | GAGATGAGACGTTGCCTCCTTC | 22 | 54.55 | 60.92 | PA | + | 1 | 0.2 |
| PA_7_RIGHT | 22 | TGCATGTGTCAGGAAGGAGTTG | 22 | 50 | 60.99 | PA | - | 1 | 0.2 |
| HA_1_LEFT | 23 | CCATGAAGACTATCATTGCTTTGAGC | 26 | 42.31 | 61.17 | HA | + | 1 | 0.4 |
| HA_1_RIGHT | 23 | AACTCCAGTGTGCCGGATGA | 20 | 55 | 61.5 | HA | - | 1 | 0.4 |
| HA_2_LEFT | 24 | TGTTACCCTTATGATGTGCCGG | 22 | 50 | 60.6 | HA | + | 2 | 0.4 |

| | | | | | | | | | |
|---|---|---|---|---|---|---|---|---|---|
| HA_2_RIGHT | 24 | CCAATAGATGCTTATTCTGCTAGGGA | 26 | 42.31 | 60.74 | HA | - | 2 | 0.4 |
| HA_3_LEFT | 25 | TTCTTTAGTAGATTAAATTGGTTGACCCA | 29 | 31.03 | 59.96 | HA | + | 1 | 0.4 |
| HA_3_RIGHT | 25 | GGAGTGATGCATTCAGACTTGC | 22 | 50 | 60.08 | HA | - | 1 | 0.4 |
| HA_4_LEFT | 26 | GGAATCTAATTGCTCCTAGGGGT | 23 | 47.83 | 59.61 | HA | + | 2 | 0.4 |
| HA_4_RIGHT | 26 | TCAATCGATTCAGCTTCCCATTGA | 24 | 41.67 | 60.95 | HA | - | 2 | 0.4 |
| HA_5_LEFT | 27 | TCATAGAAAATGGTTGGGAGGGAA | 24 | 41.67 | 60.04 | HA | + | 1 | 0.4 |
| HA_5_RIGHT | 27 | TCACATTTGTGGTATATTTTGAAACAACC | 29 | 31.03 | 60.42 | HA | - | 1 | 0.4 |
| HA_6_LEFT | 28 | GCCCTGGAGAACCAACATACAA | 22 | 50 | 60.74 | HA | + | 2 | 0.2 |
| HA_6_RIGHT | 28 | CAAGGGTGTTTTTAATTAATGCACTCA | 27 | 33.33 | 59.83 | HA | - | 2 | 0.2 |
| NP_1_LEFT | 29 | AGCAGGGTTGATAATCACTCACTG | 24 | 45.83 | 60.95 | NP | + | 2 | 0.2 |
| NP_1_RIGHT | 29 | GCGCCAGATTCGCCTTATTTCT | 22 | 50 | 61.81 | NP | - | 2 | 0.2 |
| NP_2_LEFT | 30 | TGATGAAAGAAGGAATAAATACCTGGAAG | 29 | 34.48 | 60.32 | NP | + | 1 | 0.4 |
| NP_2_RIGHT | 30 | AATTTCGATCGTTGATCCCCCG | 22 | 50 | 61.24 | NP | - | 1 | 0.4 |
| NP_3_LEFT | 31 | ATGTGCTCTCTGATGCAGGG | 20 | 55 | 59.84 | NP | + | 2 | 0.2 |
| NP_3_RIGHT | 31 | TCCCACCAAGGAATATCCCTCT | 22 | 50 | 60.21 | NP | - | 2 | 0.2 |
| NP_4_LEFT | 32 | TCACAAATCTTGCCTACCTGCC | 22 | 50 | 61.06 | NP | + | 1 | 0.2 |
| NP_4_RIGHT | 32 | GGAGGCCCTCTGTTGATTAGTG | 22 | 54.55 | 60.6 | NP | - | 1 | 0.2 |
| NP_5_LEFT | 33 | ACAAATTGCTTCAAATGAGAACATGG | 26 | 34.62 | 60.01 | NP | + | 2 | 0.4 |
| NP_5_RIGHT | 33 | TCGTACTCTTCTGCATTGTCTCC | 23 | 47.83 | 60.37 | NP | - | 2 | 0.4 |
| NA_1_LEFT | 34 | AACCAAGTGATGCTGTGTGAAC | 22 | 45.45 | 59.82 | NA | + | 1 | 0.4 |
| NA_1_RIGHT | 34 | CTGGACCATGCTATGCACACTT | 22 | 50 | 61.12 | NA | - | 1 | 0.4 |
| NA_2_LEFT | 35 | ATTCGATTAGGCTTTCCGCTGG | 22 | 50 | 61.25 | NA | + | 2 | 0.4 |
| NA_2_RIGHT | 35 | GTCATTACTACTGTACAAGTTCCATTGA | 28 | 35.71 | 59.98 | NA | - | 2 | 0.4 |
| NA_3_LEFT | 36 | ACGGGGGATGATAAAAATGCAAC | 23 | 43.48 | 60.12 | NA | + | 1 | 0.2 |
| NA_3_RIGHT | 36 | TTCTGGGTGTGTCTCCAACAAG | 22 | 50 | 60.6 | NA | - | 1 | 0.2 |
| NA_4_LEFT | 37 | AAAGGATCCAACCGGCCCAT | 20 | 55 | 61.89 | NA | + | 2 | 0.4 |
| NA_4_RIGHT | 37 | TCAGTTTCCTCTTTTCTTCCCCT | 23 | 43.48 | 59.54 | NA | - | 2 | 0.4 |
| MP_1_LEFT | 38 | GCAGGTAGATATTGAAAGATGAGCC | 25 | 44 | 59.97 | MP | + | 2 | 0.2 |
| MP_1_RIGHT | 38 | TATATGAGGCCCATGCAACTGG | 22 | 50 | 60.41 | MP | - | 2 | 0.2 |
| MP_2_LEFT | 39 | ACATGGACAAAGCAGTTAAACTGT | 24 | 37.5 | 59.68 | MP | + | 1 | 0.4 |
| MP_2_RIGHT | 39 | ACTAGGATGAGTCCCAATGGCT | 22 | 50 | 60.82 | MP | - | 1 | 0.4 |
| MP_3_LEFT | 40 | AAATGGCTGGATCAAGTGAGCA | 22 | 45.45 | 60.74 | MP | + | 2 | 0.4 |
| MP_3_RIGHT | 40 | TGCTGACAAAATGACTGTCGTC | 22 | 45.45 | 59.63 | MP | - | 2 | 0.4 |
| NS_1_LEFT | 41 | GCAGGGTGACAAAGACATAATGG | 23 | 47.83 | 60.12 | NS | + | 1 | 0.2 |
| NS_1_RIGHT | 41 | TTTCTCCATGATTGCCTGGTCC | 22 | 50 | 60.81 | NS | - | 1 | 0.2 |
| NS_2_LEFT | 42 | AGAAACTGGTTCATGCTAATGCC | 23 | 43.48 | 59.81 | NS | + | 2 | 0.4 |
| NS_2_RIGHT | 42 | TCTTCAAACTTCTGACCTAGCTGT | 24 | 41.67 | 59.98 | NS | - | 2 | 0.4 |

**Supplementary Table 4**. *RSV A Primers*. List of tiled-amplicon RSV A primers, including, name, amplicon, sequence, size, %GC, melting temperature, strand, primer pool, and pool concentration.

| Name | Amplicon | Seq | size | %gc | tm (use 65) | Strand | Pool | [pool] (uM) |
|---|---|---|---|---|---|---|---|---|
| RSV_1_LEFT | 1 | AATGCGTACAACAAACTTGCGT | 22 | 40.91 | 60.4 | + | 1 | 0.5 |
| RSV_1_RIGHT | 1 | GCCATTAGGTTGAGAGCAGTGT | 22 | 50 | 60.8 | - | 1 | 0.5 |
| RSV_2_LEFT | 2 | TGTTATTACAAGTAGTGATATTTGCCCT | 28 | 32.14 | 59.56 | + | 2 | 0.5 |
| RSV_2_RIGHT | 2 | TCATCAGTCTTTGTGGTGTGGT | 22 | 45.45 | 60.01 | - | 2 | 0.5 |
| RSV_3_LEFT | 3 | CAGAAGATAAAAATGGGGCAAATAAATCA | 29 | 31.03 | 60.12 | + | 1 | 0.5 |
| RSV_3_RIGHT | 3 | TTATGGGTGTGTGCTTGGTAGG | 22 | 50 | 60.47 | - | 1 | 0.5 |
| RSV_4_LEFT | 4 | AACACAAAATATGGCACTTTCCCT | 24 | 37.5 | 59.73 | + | 2 | 0.5 |
| RSV_4_RIGHT | 4 | GCCACATAACTTATTAATGTGTTTCTGC | 28 | 35.71 | 60.55 | - | 2 | 0.5 |
| RSV_5_LEFT | 5 | CCAGCAAATATACCATCCAACGG | 23 | 47.83 | 60 | + | 1 | 0.5 |
| RSV_5_RIGHT | 5 | GTCATGCCTGTATTCTGGAGCC | 22 | 54.55 | 61.25 | - | 1 | 0.5 |
| RSV_6_LEFT | 6 | GCAAGCTTAACAACTGAAATTCAAATCA | 28 | 32.14 | 60.8 | + | 2 | 0.5 |
| RSV_6_RIGHT | 6 | CCCTGCACCATAGGCATTCATA | 22 | 50 | 60.41 | - | 2 | 0.5 |
| RSV_7_LEFT | 7 | GGTATAGCACAATCTTCTACCAGAGG | 26 | 46.15 | 60.96 | + | 1 | 0.5 |
| RSV_7_RIGHT | 7 | ACCATTTTCTTTGAGTTGTTCAGCA | 25 | 36 | 60.49 | - | 1 | 0.5 |
| RSV_8_LEFT | 8 | TGGCCTAGGCATAATGGGAGAA | 22 | 50 | 61.16 | + | 2 | 0.5 |
| RSV_8_RIGHT | 8 | TGGTTGAATTTGATGTTATAGGGCTT | 26 | 34.62 | 59.78 | - | 2 | 0.5 |
| RSV_9_LEFT | 9 | CACATCACCCAAAGATCCCAAGA | 23 | 47.83 | 60.82 | + | 1 | 0.5 |
| RSV_9_RIGHT | 9 | ATACCATCCCGAGCAGATGTGG | 22 | 54.55 | 62.12 | - | 1 | 0.5 |
| RSV_10_LEFT | 10 | CAAACGATAATATAACAGCAAGATTAGATAGG | 32 | 31.25 | 59.98 | + | 2 | 0.5 |
| RSV_10_RIGHT | 10 | GGTGAGTTGGTTTGTTTGTTGGT | 23 | 43.48 | 60.56 | - | 2 | 0.5 |
| RSV_11_LEFT | 11 | AGTGACAATGATCTATCACTTGAAGATT | 28 | 32.14 | 59.56 | + | 1 | 0.5 |
| RSV_11_RIGHT | 11 | CCTTGGGTGTGGATATTTGTTTCAC | 25 | 44 | 60.95 | - | 1 | 0.5 |
| RSV_12_LEFT | 12 | GTGCCCATGTTCCAATCATCCA | 22 | 50 | 61.4 | + | 2 | 0.5 |
| RSV_12_RIGHT | 12 | TTTCTGACGCTGATAGATCTTAGGT | 25 | 40 | 59.79 | - | 2 | 0.5 |
| RSV_13_LEFT | 13 | TGCAGTCTAACATGCCTAAAATCAA | 25 | 36 | 59.67 | + | 1 | 0.5 |
| RSV_13_RIGHT | 13 | TTGCAAATCGTGTAGCTGTGTG | 22 | 45.45 | 60.21 | - | 1 | 0.5 |
| RSV_14_LEFT | 14 | GCCACAAAGTCAATTCATAGTAGATCT | 27 | 37.04 | 59.99 | + | 2 | 0.5 |
| RSV_14_RIGHT | 14 | CCATTGGTTGATTTTATCTAGCGTGT | 26 | 38.46 | 60.62 | - | 2 | 0.5 |
| RSV_15_LEFT | 15 | GTTCATCAGATCCAGTACTCAAATAAGT | 28 | 35.71 | 59.82 | + | 1 | 0.5 |
| RSV_15_RIGHT | 15 | GCATGGTGAGATGTTGATGTGG | 22 | 50 | 60.08 | - | 1 | 0.5 |
| RSV_16_LEFT | 16 | AGCATTCACCAATCTGATAGCTCA | 24 | 41.67 | 60.47 | + | 2 | 0.5 |
| RSV_16_RIGHT | 16 | CTTTGTGGTTTGCCGAGGCTAT | 22 | 50 | 61.64 | - | 2 | 0.5 |
| RSV_17_LEFT | 17 | AGGACCTGGGACACTCTCAATC | 22 | 54.55 | 61.34 | + | 1 | 0.5 |
| RSV_17_RIGHT | 17 | GTTGTTTTGTGGTGGGTTTGCT | 22 | 45.45 | 60.8 | - | 1 | 0.5 |
| RSV_18_LEFT | 18 | GAGTCAACCCCACAATCCACAA | 22 | 50 | 60.93 | + | 2 | 0.5 |
| RSV_18_RIGHT | 18 | GTGTTGGAGGTGAGCAGTGTAG | 22 | 54.55 | 61.05 | - | 2 | 0.5 |
| RSV_19_LEFT | 19 | ACCTCAAACCACAAAACCAAAGG | 23 | 43.48 | 60.31 | + | 1 | 0.5 |
| RSV_19_RIGHT | 19 | TGAGGATTGGCAACTCCATTGT | 22 | 45.45 | 60.41 | - | 1 | 0.5 |
| RSV_20_LEFT | 20 | TCCACTCAACCACCTCCGAA | 20 | 55 | 60.78 | + | 2 | 0.5 |
| RSV_20_RIGHT | 20 | TCTGTACCATTACACTTATTTTCCTTGAT | 29 | 31.03 | 59.81 | - | 2 | 0.5 |
| RSV_21_LEFT | 21 | GCAAAGGCTATCTTAGTGCTCTAAG | 25 | 44 | 59.91 | + | 1 | 0.5 |
| RSV_21_RIGHT | 21 | AGCCTTGTTTGTGGATAGTAGAGC | 24 | 45.83 | 60.95 | - | 1 | 0.5 |
| RSV_22_LEFT | 22 | GGCATTGCCGTATCCAAGGT | 20 | 55 | 60.77 | + | 2 | 0.5 |
| RSV_22_RIGHT | 22 | TGCTGTCTAACTATTTGAACATTGCT | 26 | 34.62 | 60.01 | - | 2 | 0.5 |
| RSV_23_LEFT | 23 | ACTACACCTGTAAGCACTTATATGTTAAC | 29 | 34.48 | 60.12 | + | 1 | 0.5 |
| RSV_23_RIGHT | 23 | TGCAGAGATTTACCTCACTTGGT | 23 | 43.48 | 59.93 | - | 1 | 0.5 |

| | | | | | | | | |
|---|---|---|---|---|---|---|---|---|
| RSV_24_LEFT | 24 | TTTTCCCACAAGCTGAAACATGT | 23 | 39.13 | 59.8 | + | 2 | 0.5 |
| RSV_24_RIGHT | 24 | TCATCAGAGGGGAACACTAATGGA | 24 | 45.83 | 61.15 | - | 2 | 0.5 |
| RSV_25_LEFT | 25 | GGGGGTGGATACTGTGTCTGTA | 22 | 54.55 | 61.08 | + | 1 | 0.5 |
| RSV_25_RIGHT | 25 | GGTGCTATTTTTATTCAGTTACTAAATGCA | 30 | 30 | 60.2 | - | 1 | 0.5 |
| RSV_26_LEFT | 26 | TATACTGCAAGGCCAGAAGCAC | 22 | 50 | 60.86 | + | 2 | 0.5 |
| RSV_26_RIGHT | 26 | CATGGGGTGGCCATTCAAAATAA | 23 | 43.48 | 60.06 | - | 2 | 0.5 |
| RSV_27_LEFT | 27 | GTCACGAAGGAATCCTTGCAAA | 22 | 45.45 | 59.63 | + | 1 | 0.5 |
| RSV_27_RIGHT | 27 | TCCTGTTGCTTTCAATATATGATATGACA | 29 | 31.03 | 59.97 | - | 1 | 0.5 |
| RSV_28_LEFT | 28 | AGAGTATGCCCTCGGTGTAGTT | 22 | 50 | 60.81 | + | 2 | 0.5 |
| RSV_28_RIGHT | 28 | GGAATTTATACTACAAGGATATTTGTCAGGT | 31 | 32.26 | 60.51 | - | 2 | 0.5 |
| RSV_29_LEFT | 29 | TGGATATCCACAAGAGCATAACCA | 24 | 41.67 | 59.91 | + | 1 | 0.5 |
| RSV_29_RIGHT | 29 | GGGATCCATTTTGTCCCATAGCT | 23 | 47.83 | 60.7 | - | 1 | 0.5 |
| RSV_30_LEFT | 30 | AATGAAATCCATTGGACCTCTCAAG | 25 | 40 | 59.73 | + | 2 | 0.5 |
| RSV_30_RIGHT | 30 | CACCTTTATGATACTTAGATATTAAGGACTGT | 32 | 31.25 | 60.02 | - | 2 | 0.5 |
| RSV_31_LEFT | 31 | TGGTCCTTATCTCAAAAATGATTATACCA | 29 | 31.03 | 59.66 | + | 1 | 0.5 |
| RSV_31_RIGHT | 31 | TGTCTGCTTTAAGATGAGATTGATTATCC | 29 | 34.48 | 60.42 | - | 1 | 0.5 |
| RSV_32_LEFT | 32 | TGGACAAGATGAAGACAACTCAGT | 24 | 41.67 | 60.22 | + | 2 | 0.5 |
| RSV_32_RIGHT | 32 | ACTAAGGCTAATATCTTTCCATGTCAAGA | 29 | 34.48 | 60.99 | - | 2 | 0.5 |
| RSV_33_LEFT | 33 | TGATAGATAATCATACTCTTAGTGGATTCCA | 31 | 32.26 | 60.37 | + | 1 | 0.5 |
| RSV_33_RIGHT | 33 | GGCATCTGTGATGTTGTTGAGC | 22 | 50 | 60.59 | - | 1 | 0.5 |
| RSV_34_LEFT | 34 | GGTTCTACATAATAAAAGAGGTAGAGGGA | 29 | 37.93 | 60.58 | + | 2 | 0.5 |
| RSV_34_RIGHT | 34 | AGCTCCTCTTAACATACTCAAACTACT | 27 | 37.04 | 60.1 | - | 2 | 0.5 |
| RSV_35_LEFT | 35 | AAGACAAGCCATGGATGCTGTT | 22 | 45.45 | 61 | + | 1 | 0.5 |
| RSV_35_RIGHT | 35 | TCTATATAATTTTGTATGTGTGACGGCA | 28 | 32.14 | 59.88 | - | 1 | 0.5 |
| RSV_36_LEFT | 36 | TCATAAATGATAAGGCTATATCACCTCCT | 29 | 34.48 | 60.27 | + | 2 | 0.5 |
| RSV_36_RIGHT | 36 | TTTCTGTAGTTCTAGATCACCATATCTTG | 29 | 34.48 | 59.51 | - | 2 | 0.5 |
| RSV_37_LEFT | 37 | TGTTCAGACAAGTTCAAATATTAGCAGAG | 29 | 34.48 | 60.92 | + | 1 | 0.5 |
| RSV_37_RIGHT | 37 | TCTATATAGTCCACTTTGCTCATCTACA | 28 | 35.71 | 59.87 | - | 1 | 0.5 |
| RSV_38_LEFT | 38 | TGTCACAATAATATGCACATATAGGCA | 27 | 33.33 | 59.61 | + | 2 | 0.5 |
| RSV_38_RIGHT | 38 | CCGTTATGTTGGATCGTTTTACTCA | 25 | 40 | 60.03 | - | 2 | 0.5 |
| RSV_39_LEFT | 39 | AGAGTATGCAGGAATAGGCCAC | 22 | 50 | 59.81 | + | 1 | 0.5 |
| RSV_39_RIGHT | 39 | GGCAAATTCATATACAATGTTAATGCTGT | 29 | 31.03 | 60.42 | - | 1 | 0.5 |
| RSV_40_LEFT | 40 | AGAGGTGAAAGTCTATTATGCAGTTTAA | 28 | 32.14 | 59.5 | + | 2 | 0.5 |
| RSV_40_RIGHT | 40 | GCATTGGGGTTTTTGTCAAACG | 22 | 45.45 | 59.89 | - | 2 | 0.5 |
| RSV_41_LEFT | 41 | CCATGATTTAAAAGATAAACTTCAAGATCTGTC | 33 | 30.3 | 60.98 | + | 1 | 0.5 |
| RSV_41_RIGHT | 41 | GCAGAAGTCTTTTCCAGTATGTTAGT | 26 | 38.46 | 59.79 | - | 1 | 0.5 |
| RSV_42_LEFT | 42 | GTTTATGAAAGTTTACCCTTTTATAAAGCAGAG | 33 | 30.3 | 61.11 | + | 2 | 0.5 |
| RSV_42_RIGHT | 42 | GGTCCTCTCTCACCACGTGTTA | 22 | 54.55 | 61.58 | - | 2 | 0.5 |
| RSV_43_LEFT | 43 | GGACATAAAATATACAACAAGCACTATAGC | 30 | 33.33 | 59.96 | + | 1 | 0.5 |
| RSV_43_RIGHT | 43 | CAAAGTGATAATTTGTAGTTCTATAAGCTGGTA | 33 | 30.3 | 61.17 | - | 1 | 0.5 |
| RSV_44_LEFT | 44 | TGGGTTAACATATGAGAAGGCCAA | 24 | 41.67 | 60.35 | + | 2 | 0.5 |
| RSV_44_RIGHT | 44 | CATATTGAGTCAAACTTATTTTGTCTGGT | 29 | 31.03 | 59.62 | - | 2 | 0.5 |
| RSV_45_LEFT | 45 | CCTCCCATATTCACAGGTGATGTT | 24 | 45.83 | 60.71 | + | 1 | 0.5 |
| RSV_45_RIGHT | 45 | ATCACACTCCAGCTTTGCTCTG | 22 | 50 | 61.06 | - | 1 | 0.5 |
| RSV_46_LEFT | 46 | GCTGGACATTGGATTCTTATTATACAACT | 29 | 34.48 | 60.62 | + | 2 | 0.5 |
| RSV_46_RIGHT | 46 | TGATAATCTATGTTAACAACCCAAGGG | 27 | 37.04 | 59.72 | - | 2 | 0.5 |
| RSV_47_LEFT | 47 | GGATGTCATAGCTTCAAACTATGGTT | 26 | 38.46 | 59.9 | + | 1 | 0.5 |
| RSV_47_RIGHT | 47 | ACCTATACAATAGTCACTCAGTGTCTT | 27 | 37.04 | 59.88 | - | 1 | 0.5 |
| RSV_48_LEFT | 48 | CACAAGTTCAATGATGAGTTTTATACTTCT | 30 | 30 | 59.71 | + | 2 | 0.5 |
| RSV_48_RIGHT | 48 | TGATGAGAAGTAGTAGTGTAAAGTTGGT | 28 | 35.71 | 60.56 | - | 2 | 0.5 |
| RSV_49_LEFT | 49 | TGTATAATTTATTTCCTACGGTTGTGATTG | 30 | 30 | 59.62 | + | 1 | 0.5 |
| RSV_49_RIGHT | 49 | ATGTTGATATGTCCATTGTACAGCC | 25 | 40 | 59.85 | - | 1 | 0.5 |
| RSV_50_LEFT | 50 | TCATAGGTGAAGGAGCAGGGAA | 22 | 50 | 60.75 | + | 2 | 0.5 |
| RSV_50_RIGHT | 50 | GTCTAATTTGAAATCGATATCATCTTGAGC | 30 | 33.33 | 60.2 | - | 2 | 0.5 |

| | | | | | | | | |
|---|---|---|---|---|---|---|---|---|
| RSV_51_LEFT | 51 | TGGAGCAAGCATGTAAGAAAATGC | 24 | 41.67 | 60.94 | + | 1 | 0.5 |
| RSV_51_RIGHT | 51 | CGTCCAGCTATAGAATATGATAGTATATCTCC | 32 | 37.5 | 61.12 | - | 1 | 0.5 |
| RSV_52_LEFT | 52 | TCCCAGTATTTAATGTAGTACAAAATGCT | 29 | 31.03 | 60.07 | + | 2 | 0.5 |
| RSV_52_RIGHT | 52 | TTTTAAGTTCATTGGTTGTCAAGCTG | 26 | 34.62 | 59.68 | - | 2 | 0.5 |

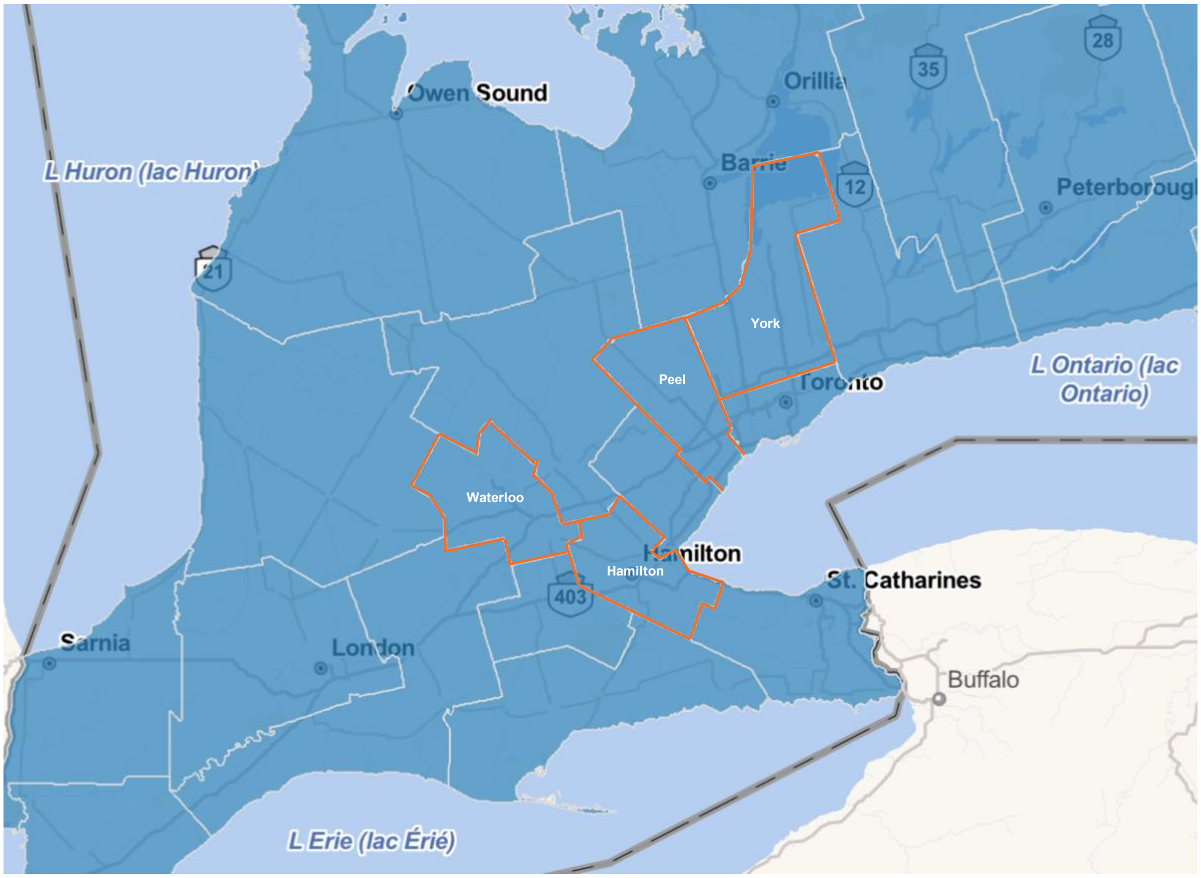


**Supplementary Figure 1**. *Location of Public Health Regions in Southern Ontario, Canada*. The boundaries of the Waterloo, Hamilton, Peel, and York public health units outlined in orange.

**Supplementary Table 5**. *List of IAV H3N2 Genes*. Segment, segment name, segment length, gene name, gene abbreviation, and location of genes of in the reference genomes, NC_007366.1 - NC_007373.1.

| Segment | Segment Name | Segment Length | Gene Name | Gene abbreviation | Location |
|---|---|---|---|---|---|
| 1 | PB2 | 2341 | Polymerase PB2 | PB2 | 28 - 2307 |
| 2 | PB1 | 2341 | Polymerase PB1 | PB1 | 25 - 2298 |
| | | | PB1-F2 protein | PB1-F2 | 119 -391 |
| 3 | PA | 2233 | polymerase PA | PA | 25 -2175 |
| | | | PA-X protein | PA-X | 25 - 784 |
| 4 | HA | 1762 | Hemagglutinin | HA | 30 - 1730 |
| 5 | NP | 1566 | Nucleocapsid protein | NP | 46 - 1542 |
| 6 | NA | 1467 | Neuraminidase | NA | 20 - 1429 |
| 7 | MP | 1027 | Matrix protein 2 | M2 | 26 - 1007 |
| | | | Matrix protein 1 | M1 | 26 - 784 |
| 8 | NS | 890 | Non-structural protein 2 | NS2 | 27 - 864 |
| | | | Non-structural protein 1 | NS1 | 27 - 719 |

**Supplementary Table 6**. *List of RSV A Genes*. Name, abbreviation, and location of genes of in the reference genome, NC_001803.1.

| Gene Name | Gene abbreviation | Location |
|---|---|---|
| Non-structural protein 1 | NS1 | 45 - 576 |
| Non-structural protein 2 | NS2 | 956 - 1097 |
| Nucleoprotein | N | 1125 - 2329 |
| Phosphoprotein | P | 2331 - 3220 |
| Matrix protein | M | 3224 - 4180 |
| Small hydrophobic protein | SH | 4190 - 4599 |
| Attachment Glycoprotein | G | 4644 - 5565 |
| Fusion protein | F | 5619 - 7521 |
| Matrix (M2/22K) | M2 | 7567 - 8527 |
| RNA-dependant RNA Polymerase | RdRp | 8460 - 15037 |
